# Parallel and Population-specific Gene Regulatory Evolution in Cold-Adapted Fly Populations

**DOI:** 10.1101/795716

**Authors:** Yuheng Huang, Justin B. Lack, Grant T. Hoppel, John E. Pool

## Abstract

Changes in gene regulation at multiple levels may comprise an important share of the molecular changes underlying adaptive evolution in nature. However, few studies have assayed within- and between-population variation in gene regulatory traits at a transcriptomic scale, and therefore inferences about the characteristics of adaptive regulatory changes have been elusive. Here, we assess quantitative trait differentiation in gene expression levels and alternative splicing (intron usage) between three closely-related pairs of natural populations of *Drosophila melanogaster* from contrasting thermal environments that reflect three separate instances of cold tolerance evolution. The cold-adapted populations were known to show population genetic evidence for parallel evolution at the SNP level, and here we find evidence for parallel expression evolution between them, with stronger parallelism at larval and adult stages than for pupae. We also implement a flexible method to estimate *cis*- versus *trans*-encoded contributions to expression or splicing differences at the adult stage. The apparent contributions of *cis-* versus *trans-*regulation to adaptive evolution vary substantially among population pairs. While two of three population pairs show a greater enrichment of *cis*-regulatory differences among adaptation candidates, *trans*-regulatory differences are more likely to be implicated in parallel expression changes between population pairs. Genes with significant *cis*-effects are enriched for signals of elevated genetic differentiation between cold- and warm-adapted populations, suggesting that they are potential targets of local adaptation. These findings expand our knowledge of adaptive gene regulatory evolution and our ability to make inferences about this important and widespread process.

## Introduction

Different species or populations often evolve similar phenotypes when adapting to similar environments (Schluter 2000; Losos, 2011). Although such parallel phenotypic evolution can be caused by amino acid changes, there is increasing evidence that regulatory mutations altering gene expression underlie many cases of phenotypic evolution (Wittkopp & Kalay, 2012; Jones et al. 2012; Stern 2013; Sackton et al. 2019). Most studies on gene regulatory evolution focus on expression abundance (the number of transcripts for a whole gene). However, alternative splicing changes resulting in modified transcript proportions can also contribute to adaptation (Barbosa-Morais et al. 2012; Gamazon and Stranger 2014; Smith et al. 2018), and yet splicing evolution has received far less study.

The level of parallelism for gene expression evolution varies across study systems. In some taxa and natural conditions, significantly more genes show parallel changes (repeatedly up- or down-regulated in one ecotype relative to the other among independent population pairs) than anti-directional changes (Zhao et al. 2015; Hart et al. 2018; Kitano et al. 2018; McGirr and Martin. 2018). However, some other cases did not show significant parallel patterns, or they even showed anti-parallel patterns (Derome et al. 2006; Lai et al. 2008; Hanson et al. 2017). The varying degree of parallelism may partly be explained by the level of divergence among ancestors: more closely related ancestors are expected to show a higher degree of parallel genetic evolution underlying similar phenotypic evolution (Conte et al. 2012; Rosenblum et al. 2014).

Furthermore, gene expression evolution can be caused by the same or different molecular underpinnings. Because of the difficulties of mapping expression quantitative trait loci (eQTLs), a first step is to classify the expression evolution into two regulatory classes. *Cis*-regulatory changes are caused by local regulatory mutations and result in allele-specific expression in a hybrid of divergent parental lines (Singer-Sam et al. 1992; Cowles et al. 2002; Yan et al. 2002; Wittkopp et al. 2004). *Trans-*regulatory changes are caused by mutations at other loci. They modify the expression of both alleles in hybrid diploids and do not result in allele-specific expression (Yvert et al. 2003; Wittkopp et al. 2004; Wang *et al*. 2007). The relative importance of *cis*- and *trans-*effects to parallel evolution varies among different studies systems (Wittkopp et al. 2008; McManus et al. 2010; Wittkopp and Kalay 2012; Coolon et al. 2014; Lemmon et al. 2014; Nandamuri et al. 2018). Many previous studies have focused on regulatory evolution between relatively distantly related lineages such as different species, from which population genetic evidence of adaptive evolution may not be available. Some studies have investigated the *cis-* vs. *trans-*regulatory variation within or between recently diverged populations but are limited to one or two populations (Chen et al. 2015; Osada et al. 2017; Glaser-Schmitt et al. 2018). To our knowledge, the only two cases comparing *cis-* and *trans*-regulatory changes for repeated adaptive divergence between populations are from threespine stickleback fish and they revealed contrasting patterns (Hart et al. 2018; Verta and Jones 2019). Hence, the relative contributions of *cis*- and *trans-*effects to recent parallel adaptation remain mostly unknown.

In part driven by interest in the evolutionary response to climate change, *Drosophila* has been used as a model system to study the genetic basis of thermal adaptation (Hoffmann et al. 2003). Because temperature is an important environmental variable along latitudinal clines, clinal populations of *Drosophila melanogaster* have been studied for decades (Adrion et al. 2015). Along these clines, populations exhibit different degrees of cold tolerance in the expected direction, suggesting spatially varying selection related to temperature (Hoffmann and Weeks 2007; Schmidt and Paaby 2008). The recent development of genomics has allowed identification of clinal genomic variants, which are candidates for thermal adaptation (e.g., Kolaczkowski et al. 2011; Fabian et al. 2012; Bozicevic et al. 2016; Mateo et al. 2018). There is also evidence of parallel evolution at the genomic and transcriptomic level (Reinhardt et al. 2014; Bergland et al. 2015; Machado et al. 2015; Zhao et al. 2015; Juneja et al. 2016; Zhao and Begun 2017). Some of these studies compared clines between species (which may have somewhat distinct biology), while others compared clines between Australia and North America (which both feature primarily European ancestry with clinally variable African admixture). Other transcriptomic studies have identified genes showing differential expression between sub-Saharan African and European populations (e.g., Catalan et al. 2012; Huylmans and Parsch 2014), which are separated by moderately strong neutral genetic differentiation associated with the out-of-Africa bottleneck.

More broadly, populations of *D. melanogaster* from contrasting environments offer an excellent opportunity to study parallel gene regulatory evolution and its underlying mechanisms. Originating from a warm sub-Saharan ancestral range (Lachaise et al. 1988; Pool et al. 2012), *D. melanogaster* has occupied diverse habitats, including environments with contrasting temperature ranges. There are at least three instances in which the species expanded to cold environments: from Africa into higher latitude regions in Eurasia, from Ethiopia lowland to higher altitudes, and from South Africa lowland to higher altitudes. Populations were collected from these six regions, representing three warm-cold population pairs: Mediterranean pair (MED), collected in Egypt (EG, warm) and France (FR, cold); Ethiopian pair (ETH) collected in Ethiopia lowland (EA, warm) and highland (EF, cold); and South Africa pair (SAF), collected in South Africa lowland (SP, warm) and highland (SD, cold). Importantly, each of these population pairs has the advantage of low genetic differentiation between its warm- and cold-adapted members compared to the differentiations among pairs (Pool et al. 2017). Although the cold populations have invaded colder habitats for only ∼1000-2000 years (∼15k-30k generations) (Sprengelmeyer et al. 2020) and different habitats have distinct selective pressures besides cold (e.g., air pressure, ultraviolet radiation, food resources), the cold-dwelling populations have shown signals of parallel adaptation for cold tolerance and allele frequency changes (Pool et al. 2017). In the present study, this unique system allows us to assess the degree of parallelism for transcriptomic changes underlying parallel adaptation to colder environments.

Because the selection environments can vary drastically across life stages of *Drosophila*, we may expect to see different patterns of local adaptation and parallelism in gene expression across stages. For *D. melanogaster*, the larvae are mostly located within fruit and their primary role is feeding. The pupae are located on or near the fruit and are immobile. The adults are mobile; their primary role is mating and reproduction (Powell, 1997; Sokolowski et al. 1986), and it is thought to be the primary overwintering stage in seasonally cold environments (Lzquierdo 1991). And a recent study using *D. melanogaster* populations across the globe found local adaptation to thermal environments at egg, larval and adult stages but not the pupal stage (Austin and Moehring 2019). Therefore, we may expect a different level of parallel gene expression evolution for thermal environments for the pupal stage.

Here, we generate RNA sequencing (RNA-seq) data for multiple outbred genotypes from each of the six population samples listed above, from larval, pupal, and adult stages. We estimate gene expression and alternative intron usage levels for each sample, then identify cases of unusually high quantitative trait differentiation between each pair of warm- and cold-adapted populations and compare their genomic locations across developmental stages. We find that genes with highly differentiated expression are enriched on the X chromosome in the adult stage relative to the larval stage. We find evidence for parallel evolution for expression for both the larval and the female adult stages, but less parallel signal for the pupal stage. We further tease out the *cis*- and *trans-*regulatory effect at the adult stage by sequencing the transcriptomes of the parental lines from different populations and their F1 offspring. Applying our resampling approach to study *cis*- and *trans-* regulatory effects, we find that the relative contributions of these effects to adaptive expression differentiation is quite variable across population pairs, with *trans*-effects showing greater parallelism. Finally, we observe enrichments of genes with high *F_ST_* among those that showed *cis-* effects and identify several candidate genes with both *cis*-effects and high *F_ST_*, as potential targets of local adaptation.

## Methods and Materials

### Ecologically and phenotypically differentiated populations

The three *Drosophila melanogaster* cold-warm population pairs used in this study, France-Egypt (MED), Ethiopia (ETH) and South Africa (SAF), were described in previous publications (Pool et al. 2012; Lack et al. 2015; Pool 2017). Previous study has shown that female adults from the cold populations (FR, EF and SD) were more likely to recover after 96 hours at 4°C than the respective warm populations (Pool et al. 2017). To extend these results, three inversion-free strains from each of the cold populations as well as an ancestral warm adapted population (ZI) were used to measure egg to adult viability at different temperatures. Viability was assayed at 15°C as the cold environment and 25°C as the warm control environment. 40 mated female flies were allowed to lay eggs in a half pint glass milk bottle with a standard medium at room temperature for 15 hours. Each strain occupied ∼8 bottles. After the flies were removed and the numbers of eggs were counted, about half of the bottles were incubated at 25°C and the other half 15°C. The numbers of adult flies that emerged from each bottle were counted after 14 days and 42 days from warm and cold environments respectively. Viability for each strain was measured as the average emergence proportion among bottles, which is the number of emerged adults divided by the number of eggs. To determine significance, unpaired t-tests between strains from each cold population and those from the ZI population were performed for both temperature conditions.

### RNA sample collection and sequencing

Within each population of the three warm/cold pairs (six populations in total), we selected 16 strains and assigned them into eight crosses (Fig. 1). Before the crossing, all the strains had been inbred for eight generations. The criterion for choosing parental strains for a cross was based on minimal genomic regions of overlapping heterozygosity. Among the strains chosen within each population, we used similar criteria to select four strains to perform crosses between the warm and the respective cold populations. Two of the four strains were used as the maternal lines and the other two were used as paternal lines in the between-population crosses. One cross between SD and SP populations was lost. We also collected adult female samples from the parental inbred lines used in the between-population crosses. Inversion frequencies are known to differ between these populations (Pool et al. 2017) and inversions have been associated with expression differences (Lavington & Kern 2017; Said et al. 2018). While inversions are not an explicit focus of our study, they may contribute to population expression differences. The inversion information for the strains used can be found in Table S1.

**Fig 1.**
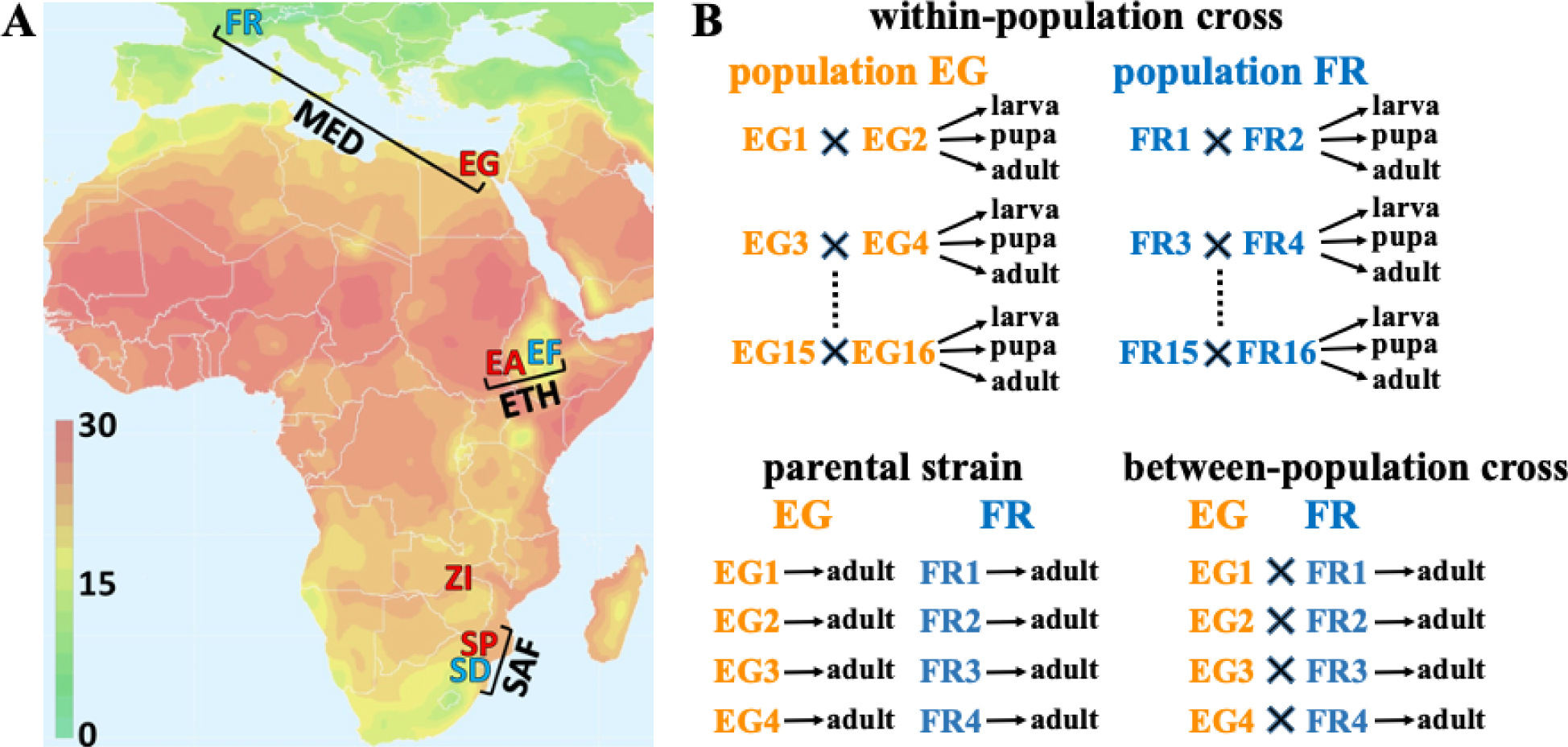
Illustrations of the geographic origins of the three population pairs and the crossing design. (A) A map of average year-round temperature (°C) showing the geographic origins of each population sample studied, and their groupings into pairs of closely-related warm- and cold-derived population samples. The Mediterranean (MED) pair comprises a cold-derived France population (FR) and a warm-derived Egypt population (EG). The Ethiopian (ETH) pair comprises a cold-derived high-altitude population (EF) and a warm-derived low-altitude population (EA). Likewise, the South African (SAF) pair comprises a cold-derived high-altitude population (SD) and a warm-derived low-altitude population (SP). The location of an additional warm-derived population from Zambia (ZI), within the species’ putative ancestral range, is also indicated. (B) Schematic figure showing the crossing design for one population pair (MED) as an example. Within-population crosses generated controlled outbred offspring for estimating *P_ST_* to quantify population differentiation in gene expression; samples at three developmental stages (third instar larva, pupa and female adult) were collected from each cross. Parental inbred strains from warm- and cold-adapted populations and inter-population crosses between them were studied to estimate *cis*- and *trans*-regulatory effects that underlie the expression divergence; samples from female adults were collected.

**Table 1.**
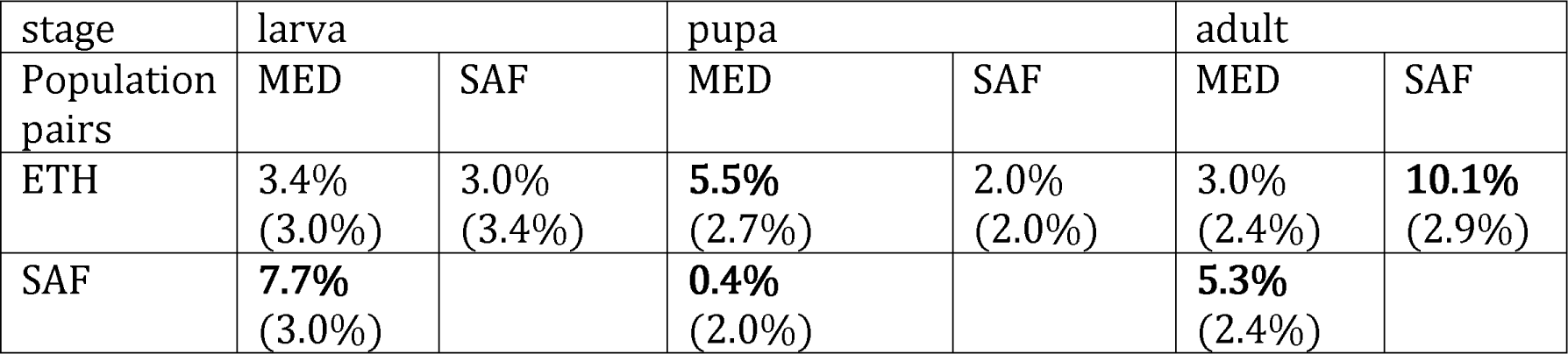
Evidence for parallel expression evolution between population pairs. The percentages of *P_ST_* outlier with parallel expression abundance changes are shown. The random expectation is the median of the permuted proportions (in brackets). The majority of proportions were higher than the expectation, with larvae and adult stages showing stronger patterns than the pupa. Those that were significantly different from the random expectation are in bold (permutation test, p < 0.01). We found one case, the MED-SAF comparison at pupal stage, that showed a significantly lower level of parallel evolution than the random expectation. Further detail regarding the numbers of shared and non-shared outliers can be found in Figure 4A.

All the flies were reared at 15°C, which approximated the derived cold condition. 20 virgin females and 20 males were collected from maternal and paternal lines respectively for each cross and allowed to mate and lay eggs for a week in half-pint bottles. Each bottle contained standard *Drosophila* medium (containing molasses, cornmeal, yeast, agar, and antimicrobial agents). For the within-population crosses, samples at three developmental stages were collected: larva, pupa and female adult. Third-instar larvae were collected on the surface of the medium. For pupa, new yellow pupae were collected within one day of pupation. For adult, female flies were collected 4-5 days after eclosion. For samples from between-population crosses and parental lines, only female adults were collected. All the samples were shock-frozen in liquid nitrogen immediately after collection.

Approximate 50 larvae or 50 pupae or 30 female adults were used for RNA extraction for each sample. Total mRNA was extracted using the Magnetic mRNA Isolation Kit (New England Biolabs, Ipswich, MA) and RNeasy MinElute Cleanup Kit (Qiagen, Hilden, Germany). Strand-specific libraries were prepared using the NEBNext mRNA Library Prep Reagent Set for Illumina. Libraries were size-selected for approximately 150 bp inserts using AMPureXP beads (Beckman Coulter, CA, USA). The 179 libraries were quantified using Bioanalyzer and manually multiplexed for sequencing. All libraries were sequenced on a HiSeq2500 (V4) with 75bp paired-end reads in two flow cells. Numbers of paired-end reads generated for each library can be found in Table S2.

**Table 2.**
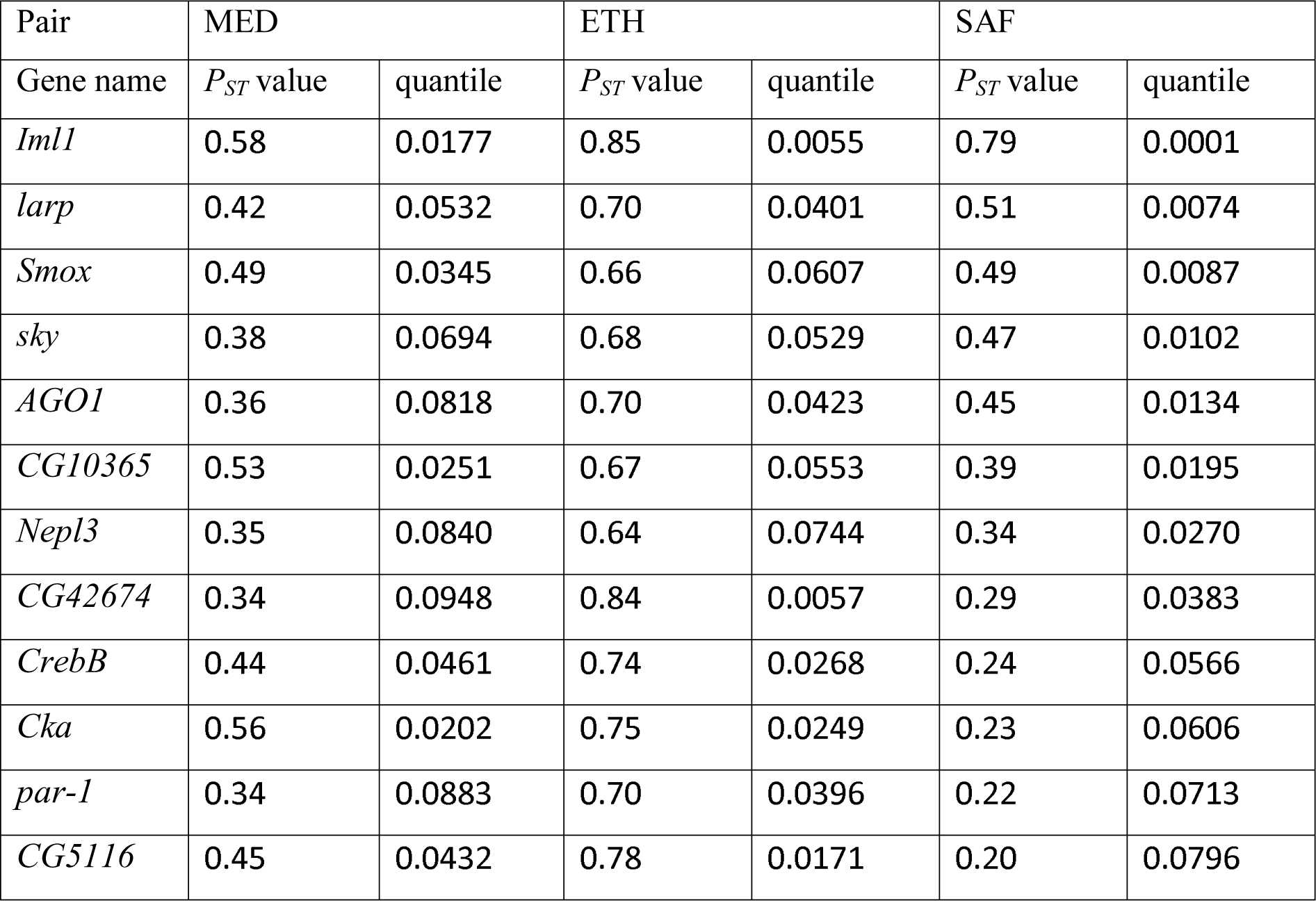
The *P_ST_* values and quantiles for the 12 genes that passed the top 10% cutoff with consistent expression changes across three population pairs at the adult stage.

### Quantifying gene expression and exon usage frequency

The paired-end sequence reads for the within-population cross samples were mapped to the transcribed regions annotated in the *D. melanogaster* reference genome (release 6, BDGP6.84) using STAR with parameters from ENCODE3’s STAR-RSEM pipeline (Li and Dewey 2011; Dobin et al. 2013). We note that cold- and warm-derived members of each population pair are expected to have very similar genome-wide reference sequence divergence (Lack et al. 2016a). For gene expression, the numbers of reads mapped to each gene were quantified using RSEM (Li and Dewey 2011). Reads mapped to the rRNA were excluded in the analysis. The expression abundance for each gene was standardized by the numbers of reads mapped to the total transcriptome of the sample.

To quantify exon usage, we used Leafcutter (Li et al. 2018) to estimate the excision frequencies of alternative introns. This phenotype summarizes different major splicing events, including skipped exons, and 5’ and 3’ alternative splice-site usage. Leafcutter took the alignment files generated by STAR as input to quantify the usage of each intron. Then Leafcutter formed clusters that contain all overlapping introns that shared a donor or accept splice site. The default parameters were used: ≥ 50 reads supporting each intron cluster and ≤ 500kb for introns length. The exon usage frequency is the number of intron excision events divided by the total events per cluster. It is worth noting that Leafcutter only detects exon-exon junction usage and it is unable to quantify 5’ and 3’ end usage and intron retention (Alasoo et al. 2019), which were not examined here.

### Principal component analysis

To visually assess the overall patterns of variation in the transcriptomes among samples, we first performed principal component analysis (PCA) for the within population cross samples across three developmental stages using *DESeq2* (Love et al. 2014). The *DESeq* dataset object was constructed from the matrix of the count data outputted from *RSEM*. After the variance stabilizing transformation (vst), the top 5000 genes with highest variance across samples at the transformed scale were used for PCA. The principal component value for each sample was obtained by the function *plotPCA* (Fig. S2). We also performed principle component analysis for samples at each developmental stage. For the adult stage, we included the F1 offspring from crosses within populations, F1 offspring from crosses between populations and the inbred parental lines of the latter crosses.

### Identifying outliers in gene expression and intron usage differentiation using within-population crosses

To identify candidate genes under differential evolution between the warm and cold populations in each pair, we first controlled for the potential transcriptome skew caused by very highly expressed genes. For each expressed gene, we calculated the average expression of the cold samples (*AvgExp_cold_*) and that of the warm samples (*AvgExp_warm_*). Then we obtained the median of the ratio of *AvgExp_cold_*/*AvgExp_warm_* across all expressed genes for the population pair. Gene expression for the warm samples was normalized by multiplying this median before subsequent analysis. This correction was designed to avoid a scenario in which either the cold population or the warm population had important expression changes in one or more highly expressed genes that caused the relative expression of all other genes to shift, even if their absolute expression level did not.

We used *P_ST_* statistics to quantify gene expression divergence between cold and warm populations in each population pair using samples from within-population crosses:

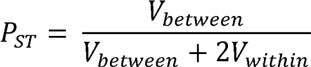

where *V_between_* is between-populations variance for expression abundance and *V_within_* is the average variance for expression abundance within populations. Although both within- and between-population components of variance can be confounded by the environmental variance, *P_ST_* is still a useful statistic to quantify phenotypic differentiation (Lande 1992; Spitze 1993; Merila 1997; Brommer 2011; Leinonen et al. 2013). Here, environmental variance should be reduced by the common laboratory environment. To reduce sampling variance before calculating *P_ST_*, for each gene, we required the total mapped reads across all 48 within-population samples to exceed 200 for a given developmental stage. Then for each population/stage, we excluded the crosses/samples with the highest and lowest gene expression for each gene (to avoid high *P_ST_* values being driven by single anomalous values), resulting in six samples per population/stage. The *P_ST_* quantile based on data excluding extreme samples is concordant with the *P_ST_* quantile calculated using all the crosses for most cases (Fig. S3).

We chose the above *P_ST_*-based approach instead of simply testing for differential expression in part because our within-population samples reflect real variation as opposed to technical replicates. Also, many alternative methods make assumptions about the data (e.g., negative binomial distribution for transcript counts) which are difficult to apply to splicing, even if they hold for expression. *P_ST_* and the population genetic index *F_ST_* are under the same theoretical framework, and are often directly compared to search for evidence of adaptive trait differentiation. However, environmental and measurement variance will downwardly bias *P_ST_*, making targets of local adaptation less likely to reach a threshold defined by genome-wide high *F_ST_* outliers. Hence, in this study we simply focus on the highest quantiles of *P_ST_* for a given trait/population comparison, as detailed below.

As with gene expression, we used *P_ST_* to estimate the intron usage differentiation between cold and warm populations, with *V_between_* as the between-population variance for a given intron’s usage frequency, *V_within_* as the average within-population variance for intron usage frequency. Before calculating *P_ST_*, for each exon-exon junction, we averaged the intron excision events (*n_i_*) and the alternative events (*n_j_*) of the cluster across all samples in a developmental stage. The minimum for both types of event had to be at least 5 (*n* ∈ [*n_i_, n_j_*] ≥ 5 We also required that at least six samples have intron usage count > 0 in each population for the exon-exon junction to be included in subsequent analysis. Then for each exon-exon junction, we excluded the sample with highest and lowest intron usage in a population/stage and calculated *P_ST_*.

### Examining the relative contribution of the X chromosome to population differentiation across stages

For gene expression differentiation, we used the upper 5% quantile of *P_ST_* as outlier cutoff to identify candidate genes potentially under geographically differential selection. Then we calculated the fraction of the outliers located on the X chromosome (*f_x_*). To generate the null distribution of *f_x_*, we permuted the genes used in calculating *P_ST_* and calculated *f_x_*’ for the top 5% of the permuted gene set. This process was repeated 10,000 times to obtain a null distribution of *f_x_*’. The upper 2.5% and lower 97.5% quantile of *f_x_*’ define the 95% confidence interval. To test whether the actual *f_x_* is significantly different from the null, the p-value is calculated as two times the proportion of the *f_x_*’ that were equal or more extreme than the actual *f_x_* (two-tailed test).

To test whether the developmental stage impacts the enrichment of outliers on the X chromosome, we analyzed the fraction of genes on the X chromosome (f) using linear model (*lm* function) in R (version 3.6.3):

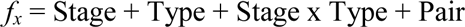

where Stage is larvae/pupal/adult. Type is outliers or nonoutliers. Pair is the population pair. As we are interested in whether the difference in *f* between outliers and the background depends on the development stage, the model above was compared to a reduced model without the interaction term Stage x Type using likelihood ratio test (*anova* function with test = “LRT”). If the Stage x Type for the full dataset was significant, we performed the same analysis separately for larva-pupa, pupa-adult and larva-adult datasets to determine which stages caused the significant Stage x Type effect.

### Comparing P_ST_ outliers with published data for African and European populations

To study whether the adult fly in MED pair changes expression in similar ways as other African and European population comparisons, we first obtained lists of candidate genes showing significant differential expression between African and European populations for adult samples from Muller et al. 2011 (padj < 0.05) and von Hecket et al. 2016. We calculated the numbers of *P_ST_* outliers that are shared with the published lists with between-populations differential gene expression (in the same directions of expression changes). Then we permuted all the genes we tested in the MED pair and selected the same number of genes as the true outliers randomly. We asked how many genes in the randomly permuted list are shared with the published lists. We repeated the process 10,000 times to obtain a null distribution of the shared numbers. The p-value was calculated as the proportion of the null distribution that was equal or more than the actual number of shared genes (one-tailed test). Another test was whether the shared genes for *P_ST_* outliers were more likely to change expression co-directionally with the published lists than the non-outliers, which was tested by a Chi-square test.

### Examining co-directional change for outliers shared between population pairs

To study the degree of parallel evolution in gene expression, we identified outlier genes shared between two population pairs and showing consistent changes in the cold populations relative to the warm ones (co-directional). Whether the number of shared outliers with co-directional change was significantly different from expected by chance was determined by a permutation-based test. For the outlier genes in a certain pair, we calculated the number of these genes (*N*) that were shared and changed expression in the same direction in the outliers from another pair. To generate the null distribution of *N*, we permuted the genes used in calculating *P_ST_* and obtained a set of genes that pass a certain quantile in each pair. Then the *N’* was calculated based on the two permuted sets of outliers. This process was repeated 10,000 times to obtain a null distribution of *N*’. To determine the statistical significance, a p-value was calculated as two times the proportion of the *N*’ that were equal or more extreme than the actual *N* (two-tailed test). The statistics here and those below assume the expression changes are independent among genes/introns, which is not always the case (genes can interact with each other via regulatory networks). We performed similar tests for pairwise comparisons between developmental stages for each population pair. The numbers for shared outliers with consistent changes between pairwise stages were reported in Table S8.

The second approach used to examine parallelism of gene expression evolution was to focus on the outlier genes for a specific population pair and examined whether the expression changes in another pair followed the same directions. If cold adaptation causes similar evolution in gene expression, those genes should tend to show changes in the same directions in both pairs. Each of the pairwise population combinations had two comparisons: the outliers can come from either pair. For the outlier genes in a certain pair, we calculated the fraction (*F*) of these genes changing expression in the same direction in another pair. To generate the null distribution of *F*, we permuted the genes used in calculating *P_ST_* and calculated *F*’ for the permuted genes that pass a certain quantile. This process was repeated 10,000 times to obtain a null distribution of *F*’. The upper 2.5% and lower 97.5% quantile of the distribution define the 95% confidence interval. The p-value was calculated as two times the proportion of the *F*’ that were equal or more extreme than the actual *F* (two-tailed test).

To identify intron usage outliers, a cutoff of the upper 5% *P_ST_* was used. If multiple exon junctions had *P_ST_* above the top 5% cutoff, only the exon junction with the highest *P_ST_* would be kept as an outlier to control for nonindependence. Because the numbers of shared intron usage outliers in both population pairs are small (<10), we only performed the second type of analysis described above. For a certain developmental stage, we used the top 5% outlier intron usage in a particular pair and asked what percentages of the intron usage changed co-directionally in another pair. To determine the statistical significance, we used the permutation approach as described above.

### GO enrichment test for P_ST_ outlier genes

The Gene Ontology enrichment tests were performed using the R package “clusterProfiler” (Yu et al. 2012) based on the fly genome annotation (Carlson 2018). The types of GO terms being tested contained all three sub-ontologies: Biological Process (BP), Cellular Component (CC) and Molecular Function (MF). Selection of overrepresented GO terms was based on adjusted p-value < 0.1 using the “BH” method (Benjamini and Hochberg 1995) for each sub-ontology. This relaxed p-value threshold (after accounting for multiple testing) was used in light of the hypothesis-generating goals of this analysis. For gene expression, the upper 5% *P_ST_* outliers were tested for GO enrichment relative to all the expressed genes for each population pair for a certain stage. To determine whether the shared significant GO terms between pairs were more than expected by chance, we randomly sampled the same numbers of genes as the outliers and performed the GO test for both pairs and identified the shared significant GO terms between pairs. We repeated the process 1000 times to get a set of numbers for the shared significant GO terms and compared to the actual number of shared significant GO terms to get a permuted p-value.

To access the functional categories of the differential intron usage, we calculated the quantile of *P_ST_* for each alternative intron’s usage. To rank the differentiation for a gene, we used the highest quantile (the most extreme differentiation) among the intron usages within the gene as the gene quantile (*q_gene_*). To account for the multiple testing of the intron usages for a gene, the adjusted total numbers of testing is calculated as 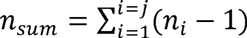, where *n_i_* is the number of testing for a cluster and *j* is the number of clusters for the gene. Then, the adjusted gene quantile is *q’_gene_* = 1- (1- *q_gene_*) x *n_sum_*. The upper 5% *q’_gene_* was used to identify the most differentiated genes for intron usage and they were tested for GO enrichment as described above.

### Estimating cis- and trans-effects of regulatory divergence using between-population crosses and parental strains

To study the contributions of *cis*- and *trans*-regulatory effects on expression and intron usage divergence, we focused our analysis on the upper 5% *P_ST_* outliers for gene expression/intron usage. For each gene/intron junction in each population pair, we selected a representative cross showing the greatest difference between parental strains for this analysis. In addition, this difference needed to be larger than the average difference between the cold and warm populations from the outbred crosses for its pair.

To study allele-specific expression/intron junction usage, we obtained the genomic sequences of the two parental strains aligned separately to the FlyBase *D. melanogaster* 5.77 assembly (Lack et al. 2015; 2016a). The SNP calling from the reference genome was done by samtools (Li et al. 2009). To avoid mapping bias for the RNAseq reads (Degner et al. 2009; Stevenson et al. 2013), we updated the reference based on the SNPs for the two parental stains by masking the SNPs as “N”. The F1 female adult RNA-seq reads were mapped to the updated reference using STAR with options: -- chimFilter None --outFilterMultimapNmax 1 (Dobin et al. 2013). Because of the fairly high level of heterozygosity within our inbred lines (Lack et al. 2015), we attempted to use polymorphic sites to study the allele-specific expression instead of focusing on the fixed difference between parental strains. However, based on the simulations we performed (see supplementary document), our new method requires a large number of F1 offspring (>300 per cross) to reduce the random sampling of parental alleles. For this experiment (only 30 F1 offspring per cross), we therefore used SNPs that were fixed differences between the parental strains. SNPs were filtered with read counts ≥ 10 in the F1 RNA-seq sample and the parental samples. Then the allele frequency in the RNA reads for the F1 sample was calculated to estimate allelic expression proportion. The allelic expression proportion for each candidate gene *p_F1_* was the median average allele frequency for all sites located in the gene region.

We tested two null hypotheses corresponding to *cis-*only and *trans*-only regulatory differences using a resampling approach. Under the null hypothesis that *cis*-regulatory effects are absent, the *p_F1_* is expected to be near 0.5 because the cold parental strain contributes half of the alleles to F1 offspring, and alleles from different parents are expressed similarly in these F1s (Cowles et al. 2002; McManus 2010; Meiklejohn et al. 2014). Under the null hypothesis that *trans*-regulatory effects are absent, *p_F1_* is expected to approximate the ratio of the cold parental strain expression to the total expression of both parental strains (Wittkopp et al. 2004): *r_F0_* = *E_c_* /*(E_c_*+ *E_w_*). However, sampling effects can cause *p_F1_* to deviate from the null expectations.

We accounted for different types of uncertainty on estimating *p_F1_*. To account for the measurement uncertainty in F1 expression, we sampled with replacement for the F1 reads mapped to each gene until we reached the numbers of reads mapped to the gene. Then we recalculated the *p_F1_’* for each SNP and then averaged across sites for each gene. We repeated the above process 1000 times to get a distribution of *p_F1_’.* A 95% confidence interval of the distribution not overlapping with 0.5 suggested the existence of a *cis*-effect.

To test for a *trans-*effect, the uncertainty when estimating the expression level in parental strains also needs to be accounted for. For each gene/intron in a parental strain, we used binomial sampling based on the expression level of the gene/intron. The sampling probability is the proportion of reads for that gene/intron relative to total reads in a sample and the number of sampling events equals the total reads of the sample. Then we had the updated expression for the cold strain *E_c_’* and the warm strain *E_w_’.* The updated *r_F0_’* is calculated as *E_c_’*/(*E_c_’*+ *E_w_’*). The sampling and calculation were repeated 1000 times. Each time the *r_F0_’* was paired with a *p_F1_’* described above to calculate the difference *D’* = *r_F0_*’ - *p_F1_’*. A 95% confidence interval of *D’* not overlapping with 0 suggested the existence of a *trans*-effect.

To test the specificity and sensitivity of this approach, we performed simulations to generate expression read data and apply our method on the simulated data (supplementary document). We found that our approach has good performance under reasonable conditions and can be adapted for other traits, such as splicing. For splicing, the *p_F1_* is the allele frequency for the diagnostic SNPs located in the exon-junction and the *r_F0_* is the ratio of the cold parental strain intron usage frequency to the sum of the frequencies for both parental strains.

Based on the tests above, the set of candidate genes were classified into categories including no significant *cis*- or *trans*-effect, cis only, and trans only (McManus 2010; Schaefke et al. 2013; Chen et al. 2015). For genes showing both *cis*- and *trans*-effects, we further classified them based on whether these two effects favored expression of the same (co-directional) or different parental allele (anti-directional). For exon usage differentiation, we applied a similar approach to classify the differentiated exons into the five categories, accounting for different sampling effects and measurement errors. Instead of analyzing expression level of the parental strains (*E*), we analyzed their intron usage frequency for the sets of outlier intron junctions.

### Examining gene expression co-regulation among outliers for adult

To study the level of co-regulation among outliers related to cold adaptation, we focused on the expressions of the outliers in the cold-derived populations. For each pairwise combination of two outliers, we calculated the correlation coefficient of the expression values among the eight outbred samples. To test whether the correlation coefficient is different from random expectation, we permuted the eight outbred samples randomly for one gene for each gene combination, requiring at least five of the eight samples to be changed. Then we calculated the correlation coefficient between genes with the permuted samples. We repeated the process 10,000 times to obtain a null distribution of the correlation coefficient. The p-value was calculated as the proportion of the null distribution that was equal or more than the actual coefficient (one-tailed test). We used p < 0.05 as a cutoff to identify significantly co-regulated outlier pairs. To compare the level of co-regulation between populations, we calculated the proportion of co-regulated pairs were significantly co-regulated for each population as well as the number of significant co-regulated partners for each gene in each population.

### Examining between-population genetic differentiation for genes with cis-effect

For the *P_ST_* outliers identified with significant *cis*-effects, we hypothesized that causative *cis*- regulatory elements may show elevated allele frequency differentiation between the warm and cold populations. For expression abundance, the majority of cis-regulatory SNPs are located within 2kb upstream of the transcription start site and downstream of the transcription end site (Massouras et al. 2012). Therefore, we used the interval from 2kb upstream to 2kb downstream as the focal region of a gene for this analysis. We calculated window *F_ST_* and SNP *F_ST_* using sequenced genomes from *Drosophila* Genome Nexus (Lack et al. 2015 & 2016a). For window *F_ST_*, the division of windows within a gene region was based on 250 non-singleton variable sites per window in the ZI population (Pool et al. 2017). Each window needed to have at least five genotypes for each population. Before assigning window *F_ST_* to the focal genes, we confirmed that there is no large chromosomal scale of differentiation between populations for each pair (Fig. S8). The highest *F_ST_* for the windows overlapping the focal region was assigned as its *F_ST_winmax_*. To determinate the statistical significance of *F_ST_winmax_*, we calculated *F_ST_winmax_* for all other blocks of the same number of windows (to account for gene length) along the same chromosome arm where cross-over rates were above 0.5cM/Mb (Comeron et al. 2012), but excluding those within 10 windows of the focal region. The specific non-low recombination regions are: 2.3–21.4LMb for the X chromosome, 0.5–17.5LMb for arm 2L, 5.2– 20.8LMb for arm 2R, 0.6–17.7LMb for arm 3L, and 6.9–26.6LMb for arm 3R. SNP *F_ST_* was calculated for sites with at least 10 alleles for each population. The highest value (*F_ST_SNPmax_*) within the focal region was thus obtained for the focal gene. Analogous to our *F_ST_winmax_* permutation, we also calculated *F_ST_SNPmax_* for permuted regions with the same number of SNPs as the focal region, along the non-low cross-over rate region on the same chromosome arm. For both *F_ST_winmax_* and *F_ST_SNPmax_*, we then focused on regions in the upper 5% quantile of permuted values for further analysis. To test whether the expression outliers with cis-effects are enriched for high *F_ST_* outliers, we analyzed the fraction of *F_ST_* outliers (*f*) using a linear model (lm) in R:

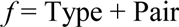

where Type is outliers with *cis*-effect (*cis*-outlier) or non-outliers, and Pair is the population pair. The model above was compared to a reduced model without the Type term using likelihood ratio test (anova function with test = “LRT”). For outlier genes that showed a *cis*-effect and high *F_ST_*, location and functional information were obtained from Flybase (Thurmond et al. 2019).

## Results

### Gene expression differentiations between warm- and cold-derived populations

In a cold environment (15°C), we found the FR and EF populations have significantly higher egg-to-adult viability than an ancestral range population and the SD population follows the same trend (Fig. S1, Table S3). In contrast, at a 25°C benign temperature all of the populations have relatively high survival (75%). These findings were consistent with past results (Pool et al. 2017) in suggesting that the cold-derived populations have adapted to low temperature.

**Table 3.**
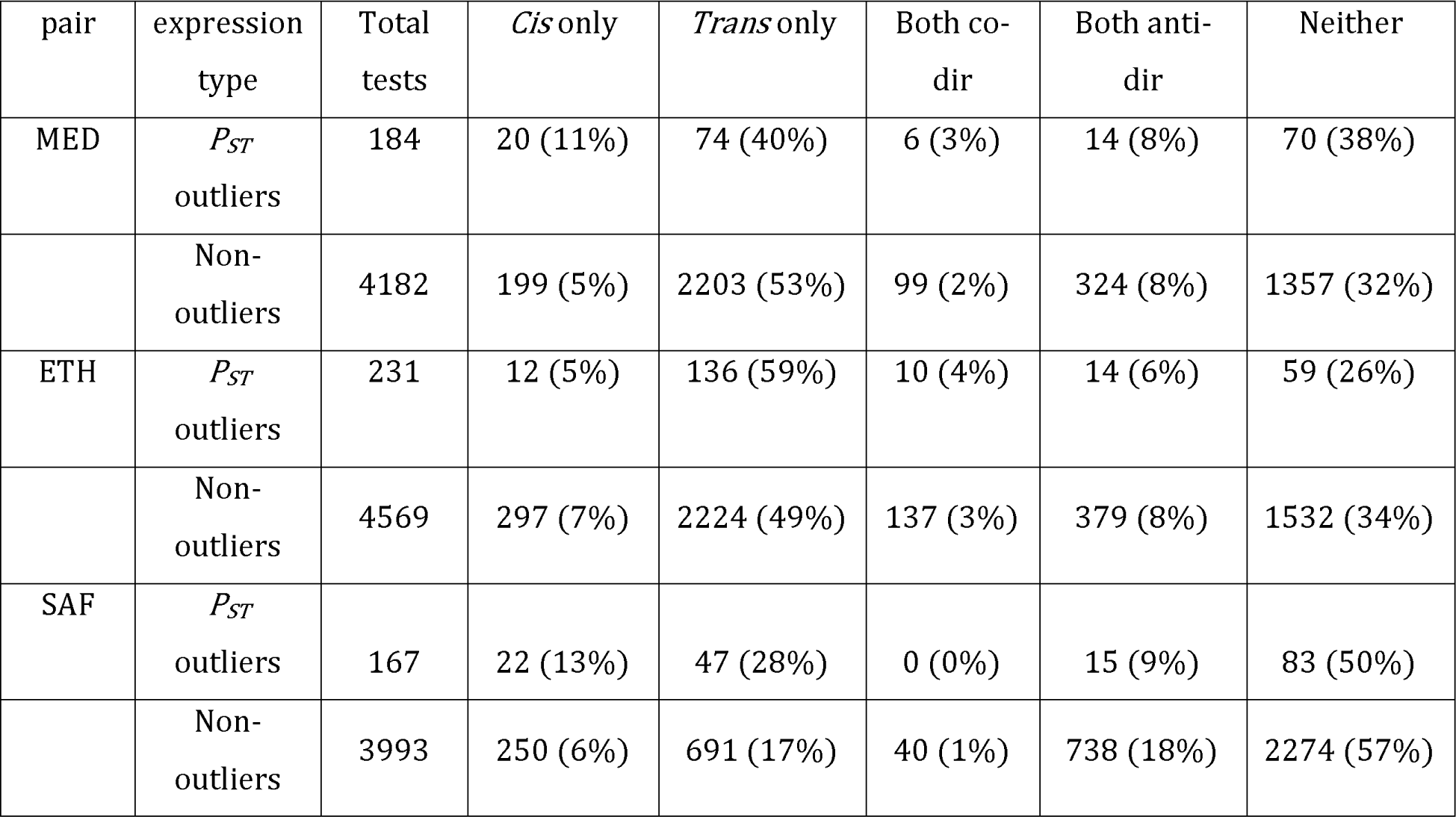
The relative prevalence of significant *cis*- and *trans*-regulatory differences for *P_ST_* outliers versus non-outliers varies among population pairs. Numbers of gene expression abundance traits showing different regulatory effects for *P_ST_* outliers and non-outliers are shown. The percentage in parentheses indicates the fraction of genes in each category relative to total genes in the tests.

We then surveyed the transcriptomes of larvae, pupae, and female adults for multiple genotypes from each cold- and warm-adapted population using high-throughput RNA sequencing (RNA-Seq). To focus on the transcriptomes of outbred genotypes, we generated eight within-population crosses from each population under the derived cold environment (15 °C). We pursued this outbred strategy to guard against inbreeding effects amplifying within-population regulatory variation and therefore dampening estimates of regulatory differentiation between populations (estimated using the quantitative genetic index *P_ST_*). Comparing our inbred adult RNA-Seq data against outbred data from a subset of the same strains, we found results that were mostly consistent with that expectation: five out of six populations showed greater expression variance in inbred than outbred data, and two out of three population pairs showed higher *P_ST_* values from outbred than inbred data (Table S6). We therefore focused on outbred data for subsequent population comparisons.

After performing PCA on normalized expression values, we found that the first and second principle component of gene expression showed clear signals of developmental stages among samples, while signals of transcriptome-wide population differentiation was more modest (Fig. S2). We then used *P_ST_* to quantify phenotypic differentiation of expression and splicing between populations in each pair. *P_ST_*, analogous to *F_ST_* for genetic variation, measures the amount of trait variance between populations versus total variance for a phenotype (Merila 1997; Brommer 2011; Leinonen et al. 2013). The genes/intron usages with highest *P_ST_* quantiles are more likely to be under ecological differential selection between populations than those with lower *P_ST_* quantiles (Leder et al. 2015).

Genes were filtered for analysis based on ≥ 200 counts across all 48 within-population samples (8 samples per population, six populations in total for three pairs). The numbers of genes that passed the filters for analysis were: 4699 genes for larva, 5098 genes for pupa and 6785 genes for adult. We initially observed that in the ETH pair, the adult sample showed a general shift in transcriptome-wide relative abundances between populations, caused by a few highly expressed genes (Fig. S4). Further investigation suggested that the EF population was primarily responsible for the observed ETH asymmetries in adults (Fig. S5). Many of the highly expressed genes in the cold-adapted and larger-bodied EF population (Lack et al. 2016b) are related to muscle protein (Table S4). To correct for the influence of such changes on relative expression levels, we standardized the expression values of warm-derived populations by the median expression ratio between cold- and warm-derived populations, resulting in about the same numbers of genes with increased and decreased expression in the cold-derived population relative to the warm-derived one transcriptome-wide (Fig. S4). To study gene expression divergence potentially under ecologically differential selection, we calculated *P_ST_* (Materials and Methods). The *P_ST_* values for all genes for each population/stage are listed in Table S4. We used the upper 5% quantile of *P_ST_* as outliers for each population pair. For the outliers, there is a strong directionality on the expression difference between populations for the ETH pair (Fig. 2): a large majority of ETH *P_ST_* outliers had higher expression in the cold-adapted EF population in larvae and especially adults, with pupae showing a reversed pattern. These asymmetries mirrored transcriptome-wide skews in expression proportion for this population pair, in that substantially more genes had higher EF expression than the modal proportion (Fig S4). These observations suggest unique regulatory features for the populations in the ETH pair, perhaps hinting that many outlier genes might be co-regulated.

**Figure 2.**
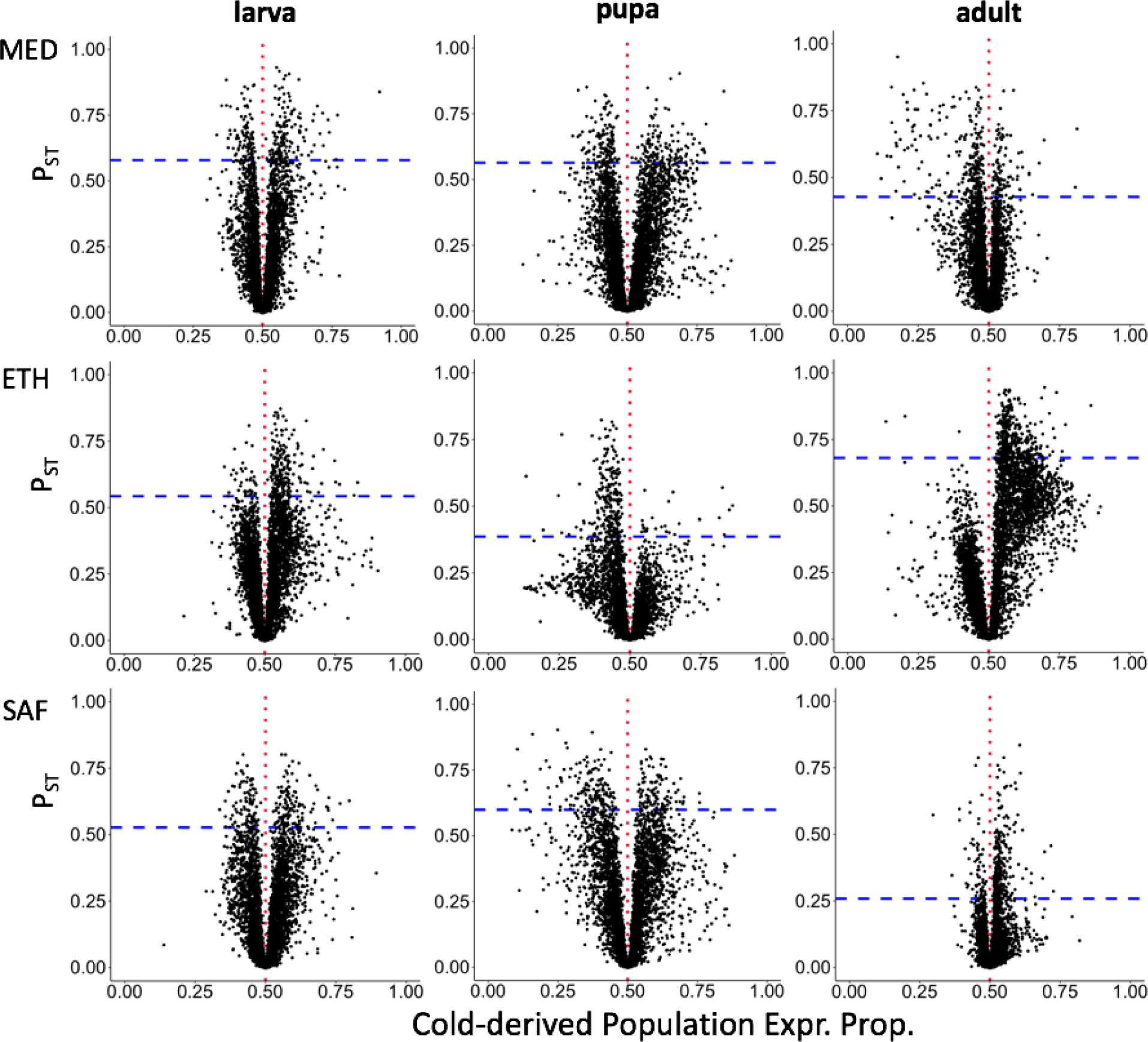
The relationship between expression differences between populations and *P_ST_* illustrates asymmetric expression differences in some population pairs / developmental stages. The x-axis is the cold-derived population expression proportion, which is the ratio of mean expression of the cold-derived population relative to the sum of mean expression values from the two populations. Proportion higher than 0.5 (red vertical dashed lines) indicate a higher expression for the cold-derived population and the warm-derived one. The blue horizontal dashed lines show the upper 5% quantile of *P_ST_*.

### Comparing P_ST_ outliers with published studies

Since there are studies comparing whole-body transcriptomes between African and European populations in this species (Muller et al. 2011; von Heckel et al. 2016), we examined whether our *P_ST_* outliers for adults from the MED pair overlapped with the candidates showing differential expression between African and European populations from the published datasets. While each of these analyses might detect regulatory evolution that occurred in Europe, we emphasize the distinctness of these geographic comparisons, since our Egypt sample differs substantially from the Zimbabwe population featured in those studies. For upper 5% quantile of *P_ST_* outliers, we do not see more sharing with the previous candidates from either study than random expectations (permutation test, p = 0.47 for comparing with Muller et al. 2011 and p = 0.39 for comparing with von Heckel et al. 2016). However, for the upper 10% of *P_ST_*, the outliers of MED are more likely to be shared with the candidates from Muller et al. 2011 than random expectations (permutation test, p = 0.0033) but not with those from von Heckel et al. 2016 (permutation test, p = 0.47). Moreover, among all the shared genes, our outliers were more likely to show gene expression change in the same directions as the previous candidates than the non-outliers (χ^2^ = 6.3, df = 1, p = 0.012 for comparing with Muller et al. 2011, χ^2^ = 7.4, df = 1, p = 0.0065 for comparing with von Heckel et al. 2016).

### X chromosomal and autosomal contributions to regulatory evolution

To investigate the contribution of autosomes versus the X chromosome to the expression differentiation due to cold adaptation, we surveyed the locations of *P_ST_* outliers for different developmental stages (Fig. 3). At the larval stage, the proportions of *P_ST_* outliers located on the X chromosome were lower than the genome-wide level in each population pair (permutation test, p = 0.72 for MED; p = 0.0076 for ETH; p = 0.021 for SAF). In contrast, at the adult (female) stage, the proportions of outliers located on the X tended to be higher than the background level (permutation test, p = 0.30 for MED; p = 0.013 for ETH and p = 0.89 for SAF). For pupa, the X chromosome enrichment was not significantly different between outliers and the background. Considering the three population pairs together, the relative enrichments of outliers on the X chromosome were significantly different among developmental stages (likelihood ratio test, p = 0.048). The patterns of different enrichment among stages were caused by larva/adult differences (likelihood ratio test, p = 0.0058 for larva-adult; p = 0.25 for pupa-adult; p = 0.24 for larva-pupa). The larva/adult difference might suggest either (1) a greater abundance of genes affecting adult fitness on the X chromosome, as previously suggested (Gibson et al. 2001; Innocenti and Morrow 2010), (2) a greater influence of female fitness on X chromosome evolution (e.g. Vicoso & Charlesworth 2006), in light of sex differences between our adult (female) and larval (mixed sex) samples, or (3) differences in the contributions of local adaptation and genetic drift to outlier sets at different stages, in combination with differential effects of drift between the X chromosome and autosomes (e.g. Pool & Nielsen 2007).

**Fig 3.**
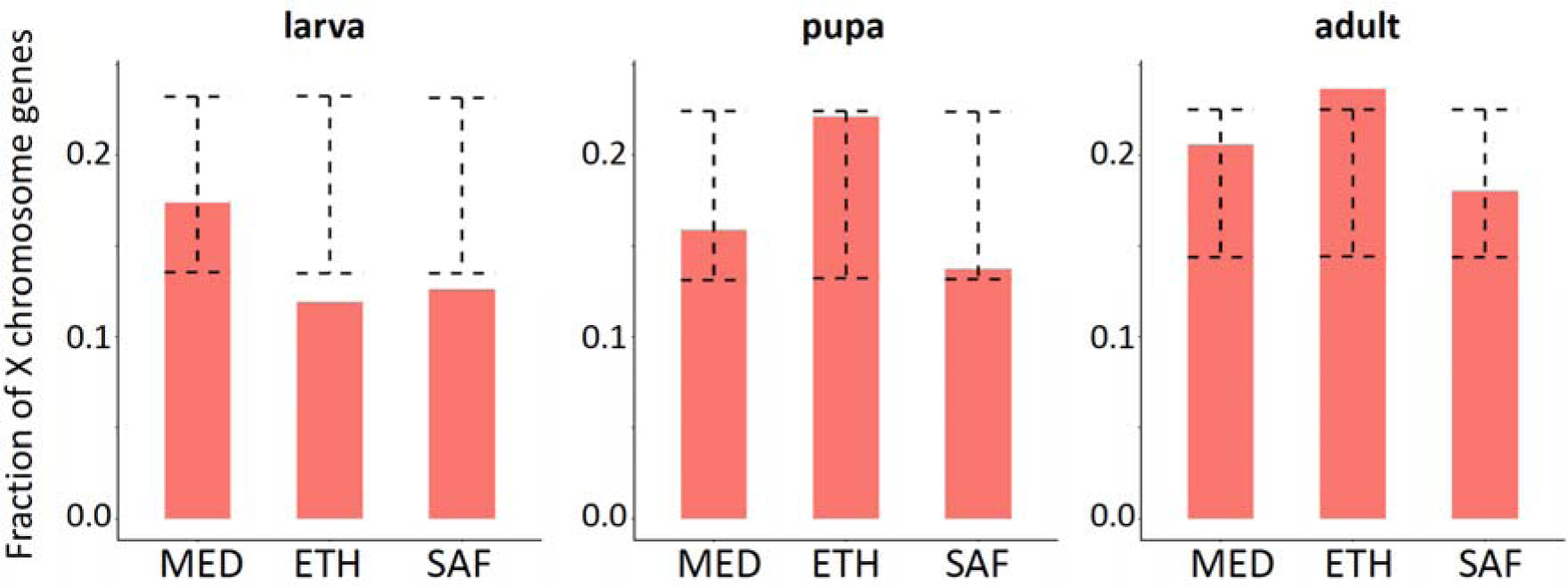
The fraction of *P_ST_* outliers located on the X chromosome varies between developmental stages. The dashed error bar indicates the 95% confidence interval for the permuted data. If the real data (orange bar) is outside the range of the error bar, it indicates the fraction is significantly different from the genomic background (p < 0.05).

### Co-directional evolution in gene expression between population pairs

Among the upper 5% of *P_ST_* outliers, we found at least some significant signals of parallel expression divergence in all three pairwise comparisons (MED vs. ETH; MED vs. SAF; ETH vs. SAF), where the shared outliers with co-directional changes (i.e., expression difference between cold- and warm-derived populations in the same direction for both pairs) were more abundant than expected by random permutation (Table 1; Fig 4A). Averaged across three pairwise comparisons, 6.7% of the outliers were shared and changed consistently for the adult stage while 4.7% of outliers were co-directional for larva and only 2.6% were co-directional for pupa (for which one population pair comparison for pupa showed significant anti-directional changes). Changing the *P_ST_* outlier cutoffs to 2.5% or 10% produced qualitatively similar patterns (Table S7). We found one shared outlier with co-directional changes among all three pairs for the adult stage: *Iml1*, a regulator of cell size, starvation response, TOR signalling, and meiosis initiation (Bar-Peled et al. 2013; Wei et al. 2014). No three-pair shared outliers were found for the larva and pupa stages. One shared outlier among three pairs is not significantly more than the expectation by permutation. Still, it is worth noting that changing the *P_ST_* outlier cutoff to 10% results in 12 shared outliers with co-directional changes among three pairs (permutation test, p < 0.0001), which suggests that meaningful regulatory evolution may extend beyond our defined outliers. The names, *P_ST_* values and quantiles of these 12 genes are shown in Table 2. Of these genes, we note the role of *CrebB* in circadian behavior, which is known to differ between *D. melanogaster* populations from different thermal environments (Svetec et al. 2015; Cao & Edery 2017; Helfrich-Förster et al. 2018). Overall, these analyses suggest that the adult stage has the highest level of parallel evolution while pupa has the lowest.

**Fig. 4.**
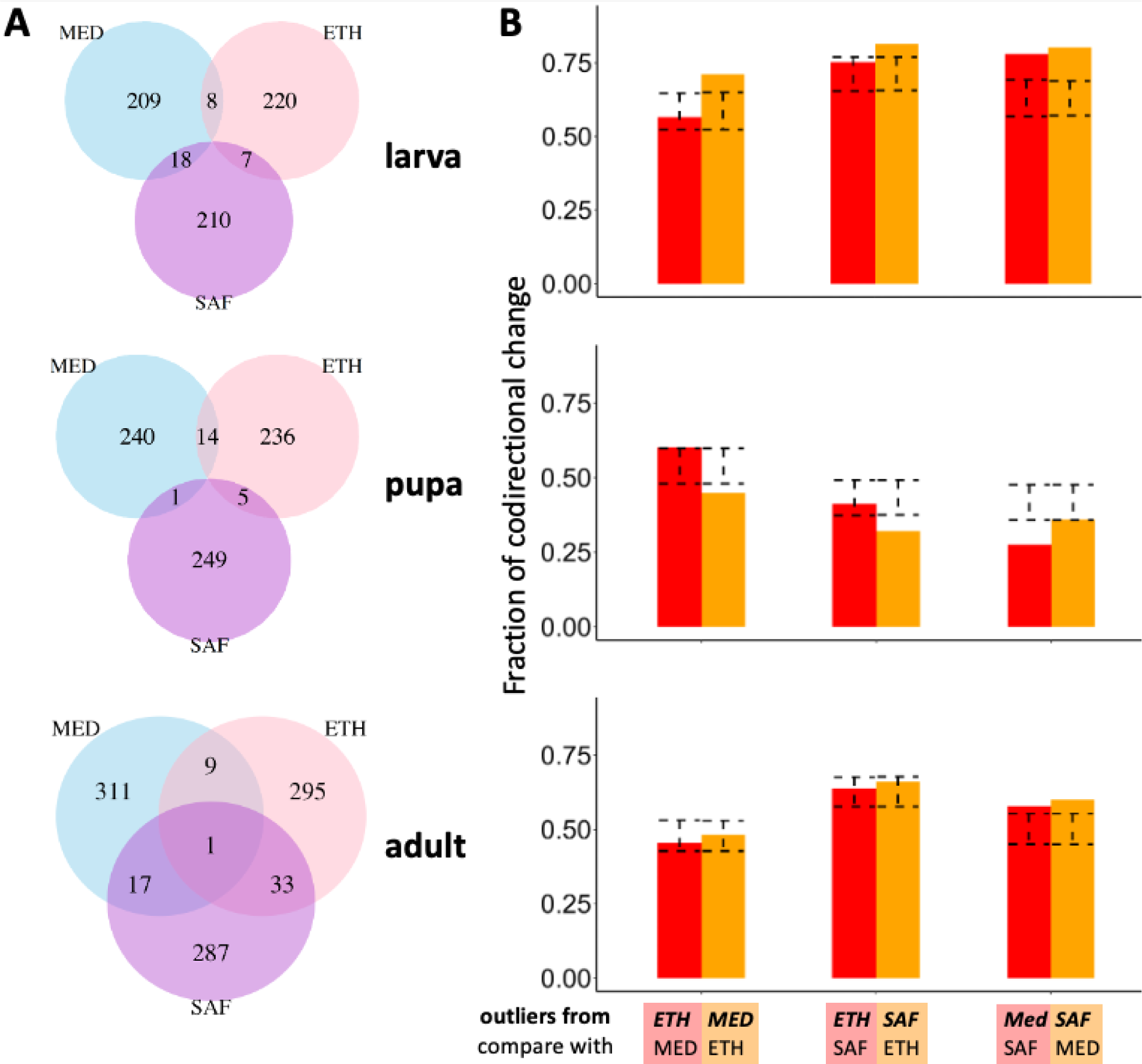
Extent and direction of parallel evolution in gene expression among population pairs. (A) Venn diagrams for the numbers of shared outliers with co-directional changes between pairs and the rest of genes. (B) Fractions of co-directional gene expression changes between pairs for the *P_ST_* outliers identified in one pair of them. The name above is the pair used to identify outliers. The name below is the other pair in the comparison. The outliers used for each bar is the summed number of the pair in the Venn diagram. The dashed error bar indicates the 95% confidence interval for the fraction of co-directional expression changes in permuted data. If the real data is outside the range of the error bar, it indicates the fraction is significantly different from random expectation (p < 0.05, two-sided test based on permuted distribution).

The analysis above requires genes being outliers in both population pairs, which is quite restrictive (the naïve expected proportion of sharing between two pairs under a 5% *P_ST_* cutoff is 5%x5%x0.5 = 0.125%) and may miss some broader patterns of parallel changes. We therefore performed a complementary analysis which only required genes being outliers in one population pair and examined whether the expression for this set of genes changed in the same direction in another pair, regardless of outlier status in the latter pair. For example, 235 genes were outliers in the MED pair at the larval stage. Then in another pair, we calculated the fraction of the 235 genes with expression differences between cold- and warm-derived populations in the same direction as for MED. There were excesses of co-directional changes for the larval stage (Figure 4B). The patterns were weaker for the adult stage and there were excesses of anti-directional changes for the pupal stage. Changing the *P_ST_* cutoff to 2.5% or 10% produce qualitatively similar patterns (Fig. S6).

We also performed similar analyses for *P_ST_* outliers of alternative intron usage. The numbers of intron-excision junctions that passed the cutoffs for *P_ST_* calculation were 4520 for larva, 5574 for pupa, and 7367 for adult. The adjusted gene quantiles for splicing are listed in Table S5. The patterns of co-directional changes were qualitatively similar to those for gene expression (Fig. S7). The fractions of co-directional changes were still highest for the larvae among the three stages; all of the comparisons except one showed an excess of co-directional changes relative to the background level, although only one comparison is significant based on the permutation test (outliers from MED being co-directionally expressed in SAF). Overall, the patterns for co-directional changes were weaker for splicing than those for gene expression. Table S9 lists the genes with the top 20 *P_ST_* for both expression and intron usage for each population pair. An interesting example is *curled*, which is an extreme splicing outlier for ETH and MED, and an extreme expression outlier for ETH as well. Also known as *nocturnin*, one isoform of this gene is thought to have a dedicated role in circadian regulation (Nagoshi et al. 2010).

### Enriched functional categories for the P_ST_ outliers

Significant Gene Ontology (GO) terms enriched in different sets of *P_ST_* outliers for gene expression are listed in Table S10. Among the significant GO terms for different population pairs, we found five terms shared between the MED pair (24 significant terms) and ETH pair (47 significant terms) at the adult stage (mitochondrion, nucleoside metabolic process, ribonucleoside metabolic process, purine nucleoside metabolic process and oxidation-reduction process). The level of sharing was significantly more than expected by chance based on permuted outlier sets (p < 0.001, no shared GO terms were found in the permuted datasets), suggesting functional convergence for adult development to the cold environment for the MED pair and ETH pair. Further, similar (though non-identical) GO terms were identified from different pairs at different stages such as terms related to mitochondria, nucleoside metabolism, and oxidoreductase complex. However, the majority of GO terms were unique for different pairs, suggesting that many functional changes for adaptation to cold environments may be population-specific. For intron usage, we only found one significant GO term for SAF pair at the larval stage (meiotic cell cycle).

### Cis- and trans-acting contributions to differential gene expression at the adult stage

Gene expression differentiation can be caused by *cis*- or *trans-*regulatory effects. A *cis*-effect comes from a local regulatory mutation and results in an allele-specific expression difference in a F1 hybrid; while a *trans-*effect is caused by remote loci that modify both alleles in a diploid cell. Therefore, a *cis*-effect can be estimated by the allelic expression in F1 hybrid (Cowles et al. 2002). We quantify that effect based on the expression proportion in the F1 offspring of a between-population cross for the allele from the cold population minus 0.5 (null expectation when *cis*-effect absent). A *trans-*effect can be estimated by the expression difference between parents that was not attributed to the *cis-*effect (Wittkopp et al. 2004), as described in the Materials and Methods.

First, we described the transcriptome-wide patterns of *cis*- and *trans-*effect sizes across all analyzed genes at the adult stage. The magnitudes of *trans-*effect sizes were significantly larger than the *cis*-effect sizes in all three population pairs (mean absolute *cis*-effects and *trans*-effects were: MED pair, 0.07 vs. 0.16, p < 2.2e-16; ETH pair, 0.07 vs. 0.16, p < 2.2e-16; SAF pair, 0.09 vs. 0.11, p < 2.2e-16. ‘Mann-Whitney’ paired test.). Moreover, we found strong negative relationships between *cis*- and *trans-*effects within each population pair (Fig. S7), where the *cis*- and *trans-*effects were generally in the opposite directions. Although the pattern can be biologically meaningful, it may also represent an artifact from using the same F1 expression data for allele specific expression (ASE) estimation to infer both *cis*- and *trans*-effects. Any measurement error on ASE will introduce an artifactual negative correlation between *cis*- and *trans*-acting changes (see Discussion below).

Next, we used our flexible permutation approach (see Materials and Methods and supplementary document) to study how many genes show a significant *cis*-effect, *trans-*effect, or both among the three population pairs (Table 3). Fig. 5A shows a graphical depiction of *cis-* and *trans-*effects on F1 and parental samples, illustrating *cis*-effects influence both F1 allele-specific expression and the parental expression ratio while *trans*-effects only influence the parental expression ratio. Averaged across population pairs, about 60% of genes show significant *cis*- and/or *trans-*effects (62% for outliers and 59% for non-outliers). We also found that 19% of genes show *cis*-regulatory effects while 53% of them show *trans-*effects, consistent with *trans-*effects being stronger on average than *cis*-effects. This apparently greater prevalence of *trans-*regulatory evolution was observed in spite of our lesser power to detect *trans-* relative to *cis*-effects (Supplementary Text).

**Fig 5.**
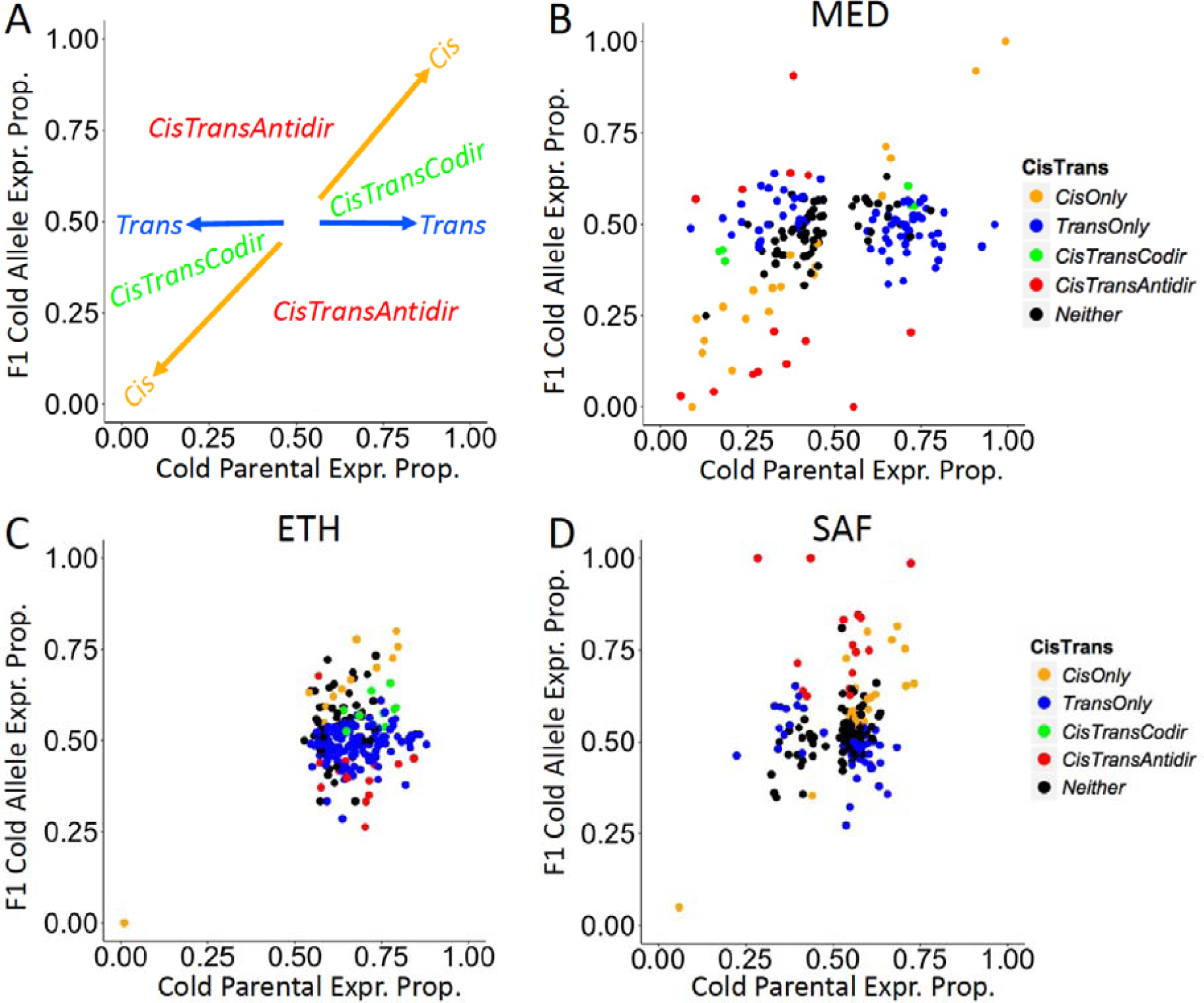
Population pairs show distinct patterns of *cis*- and *trans*-regulatory evolution for expression outliers. These plots depict evidence for *cis-* and *trans*-regulatory evolution based on the relative expression proportion of cold strain alleles in parental and F1 datasets. (A) Conceptual depiction of *cis*- and *trans*-effects. *Cis-*effects change the F1 allele-specific expression and the parental expression proportion concordantly (along the one-to-one ratio line). *Trans-*effects only change the parental expression proportion but not the F1 allele-specific expression (vertical line y = 0.5). Co-directional *cis-* and *trans-*effects (*CisTransCodir*) locate in the two spaces within the 45° angle between the *Cis* vector and the *Trans* vector. Anti-directional *cis*- and *trans*-effects (*CisTransAntidir*) locate in the two spaces within 135° angle between *Cis* vector and *Trans* vector. (B-D) Evidence for *cis-* and *trans*-regulatory evolution of putatively adaptive expression differences between warm- and cold-derived populations (expression *P_ST_* outliers).

To examine the *cis*- and *trans-*regulatory contributions to adaptive evolution of gene expression, we compared genes showing *cis-* or *trans*-effects between *P_ST_* outliers (Fig. 5B-D) and non-outliers. Because of the potential artifact generating opposing *cis*- and *trans-*effects, we excluded genes showing both significant *cis*- and *trans-*effects in opposite directions (Both anti-dir category in Table 3). For genes showing significant *cis*-effects (Fig. 6A), they were enriched in the outliers relative to the non-outliers for the MED pair (χ2 = 11.6, df = 1, p = 0.00066) and the SAF pair (χ2 = 4.8, df = 1, p = 0.029) but not for the ETH pair (χ2 < 0.01, df = 1, p = 1). While for significant *trans*-effects genes (Fig. 6B), they were enriched in the outliers relative to the non-outliers for the ETH pair (χ2 = 9.6, df = 1, p = 0.0019) and the SAF pair (χ2 = 5.4, df = 1, p = 0.020) but the enrichment is opposite for the MED pair (χ2 = 10.2, df = 1, p = 0.0014).

**Fig 6.**
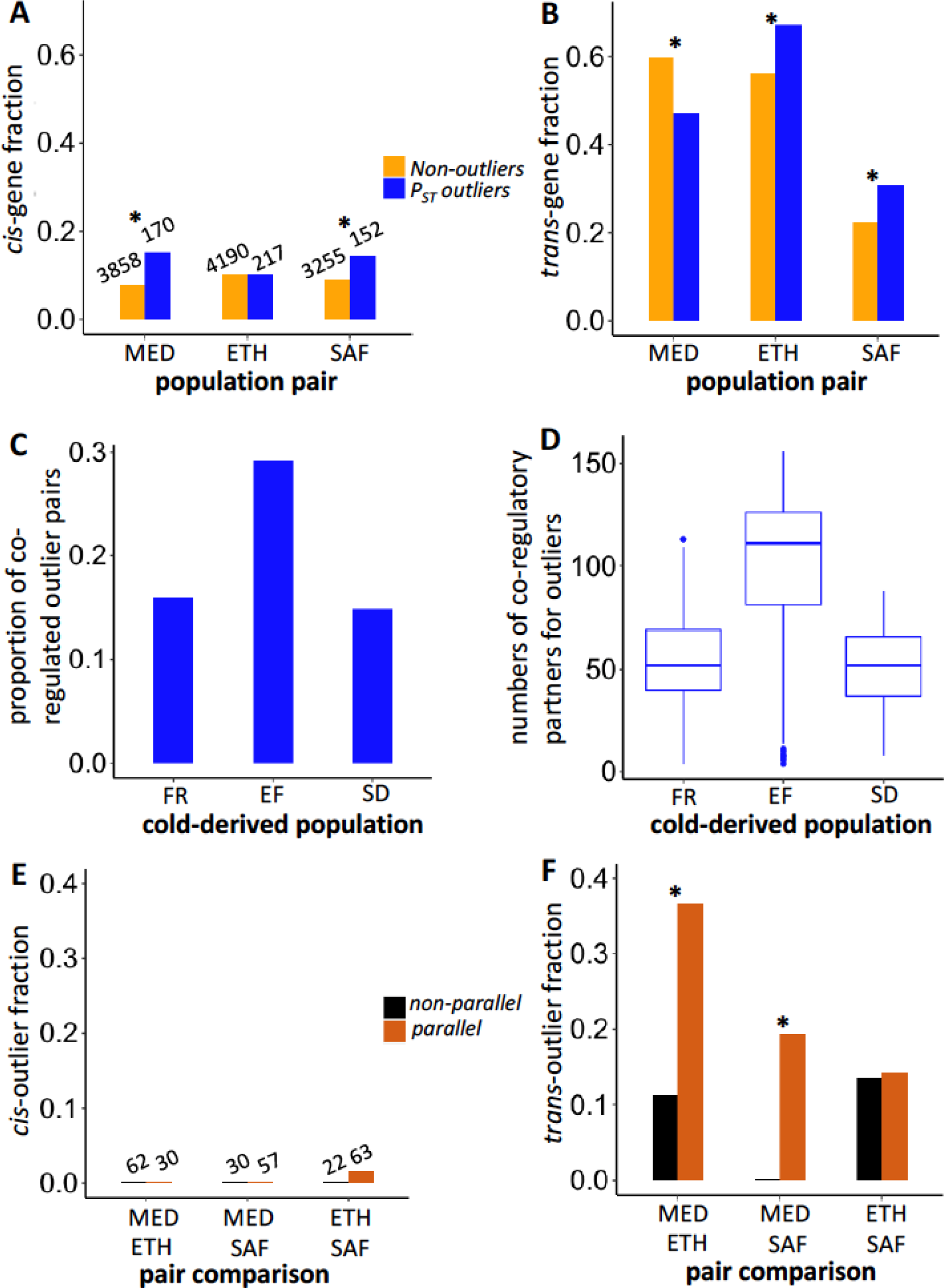
The prevalence of *cis*- versus *trans*-regulatory *P_ST_* outliers differs between population pairs, while trans changes show greater parallelism. The upper panels show the fractions of genes with *cis* regulatory effects (A) and those with *trans*-effects (B) for *P_ST_* outliers and non-outliers in each population pair. The middle panels show the proportion of co-regulated pairs among all pairwise outliers (C) and the numbers of co-regulatory partners for outlier genes (D) in each cold-derived population. The lower panels show the fractions of parallel outlier genes (*i.e.* shared and codirectional between population pairs) and non-paralleled outlier genes (*i.e.* shared and anti-directional between population pairs) showing *cis*-effect (E) and *trans-*effect (F) in different population pairs. The number above the bar for the *cis*-effect shows the denominator of the fraction, which applies to the *trans-*effect results as well. * indicates the fractions are significantly different between two categories (p < 0.05).

Because theory suggests that co-regulation of genes can amplify the contribution of *trans*-effects (Liu et al. 2019), we examined the level of co-regulation between pairwise outliers by calculating the correlation coefficient between the expression values among the eight outbred samples within each cold-derived population. Indeed, the percentage of pairwise correlation with evidence of co-regulation (p-value < 0.05) is much higher in EF population than that in FR and SD populations (Fig. 6C, Table S11). At the gene level, the median number of significant co-regulatory partners for EF is about twofold that for FR and SD (Fig. 6D). This pattern supports the hypothesis that substantial co-regulation of outlier genes results in more significant *trans*-effects for the ETH pair (Fig. 6B). Further, the strong co-regulation in EF might be related to the asymmetry observed in the ETH outliers (Fig. 2). However, the pattern could reflect variation in cell type content among strains rather than co-regulation within cells (see Discussion).

To study the role of *cis*- and *trans-*regulatory effects on the parallel adaptation, we further focused on the outlier genes. We categorized the top 10% *P_ST_* outliers based on whether they were shared between two pairs with consistent expression changes (parallel) or opposite changes (non-parallel). Because genes with *cis*- (or *trans-*) effects in both population pairs in the parallel category would indicate the contribution of *cis*- (or *trans-*) effects to parallel adaptive evolution, we identified shared outlier genes with *cis*- (or *trans-*) effects in any two pairs in both parallel and non-parallel categories (excluding genes showing both significant *cis*- and *trans*-effects in opposite directions). For *cis*-effects, only one gene (*CG42788*) is significant in both ETH and SAF pairs (Fig. 6E). While for *trans-*effects, the numbers of genes showing *trans*-effect are four for the MED and ETH pairs (*CG8034*, *CG33981*, *kst* and *wdb*), two for the MED and SAF pairs (*scf* and *AP-2*) and four for the ETH and SAF pairs (*CG7766*, *Dlg5*, *larp* and *Pink1*). Of these, *larp* had *P_ST_* quantiles near or below 0.05 in all three population pairs (Table 2); its functions include mitochondrial regulation (Zhang et al. 2019). There are significant enrichments of *trans*-effect genes in the parallel outlier category relative to the non-parallel category in two out of three population pair comparisons (Fig. 6F, MED & ETH: p = 0.0097; MED & SAF: p = 0.014; ETH & SAF: p = 1; Fisher’s Exact Tests). As a complementary analysis, we also included the non-shared outliers in the non-parallel category, and we found qualitatively similar patterns for *cis*- and *trans*-effects (Fig. S9). Overall, there is stronger evidence of *trans-*regulatory evolution contributing to parallel gene expression changes between cold-adapted populations.

### Cis- and trans-acting contributions to differential intron usage at the adult stage

For all intron usage traits, we found the magnitude of *trans*-effects on average to be higher than that of *cis*-effects (mean absolute *cis*-effects and *trans*-effects are: MED pair, 0.13 vs. 0.17, p = 0.0001; ETH pair, 0.30 vs. 0.32, p = 0.04; SAF pair, 0.17 vs. 0.19, p = 0.0044. ‘Mann-Whitney’ paired test). Because of the limited diagnostic SNPs with enough read depth located in the intron junction regions, there are few outlier introns tested for *cis*- and *trans*-regulatory effects (Table S12).

### Elevated genetic differentiation at cis-regulated expression outliers

Since the *cis*-regulatory mutations contributing to local adaptation may show differentiation in allele frequency between populations, we examined genetic differentiation for expression outliers with *cis*-effects (including *cis*-only genes and genes with both *cis*- and *trans*-effects to increase sample sizes). We examined whether each of these *cis*-outliers shows high *F_ST_* between that pair of cold- and warm-adapted populations – for both window *F_ST_* (*F_ST_winmax_*) and maximum SNP *F_ST_* (*F_ST_SNPmax_*). A gene showing both significant *cis*-effect and higher *F_ST_* quantile (here, the top 5% versus comparable genomic regions) could reflect adaptive regulatory evolution targeting the surveyed sequences or nearby sites. We first confirmed that there is little chromosomal scale differentiation between populations (aside from known moderate X-autosome differences for the MED pair; Lack et al. 2015) by plotting window *F_ST_* across the major chromosome arms (Fig. S8). We observed no obvious clustering of *P_ST_* outliers along the chromosome arms. For window *F_ST_* (Fig. 7A), high *F_ST_* is enriched in *cis*-effect outliers relative to the non-outliers for the MED pair (p = 0.022. Fisher’s Exact Test) and the other two pairs show the same trend (ETH: p = 0.22; SAF: p = 0.23). Although the MED result would be only marginally significant if correcting for the three tests performed, a clearer result is obtained when considering the three population pairs together. In this analysis, the fractions of genes with high window *F_ST_* were significantly higher in *cis*-outliers than that in non-outliers (likelihood ratio test, p = 0.0056). For maximum SNP *F_ST_* (Fig. 7B), the MED and SAF pairs showed significant enrichments of high *F_ST_* in *cis*-effect outliers versus the non-outliers, while the ETH pair showed a weak trend in that direction (MED: p = 0.012; ETH: p = 0.49; SAF: p = 0.030). Similar analysis combining the three pairs found that the fractions of high maximum SNP *F_ST_* were significantly higher in *cis*-outliers than that in non-outliers (likelihood ratio test, p = 0.0045). These results reflected more than two-fold enrichment of *F_ST_* outliers among *cis*-regulated *P_ST_* outliers across population pairs.

**Fig. 7.**
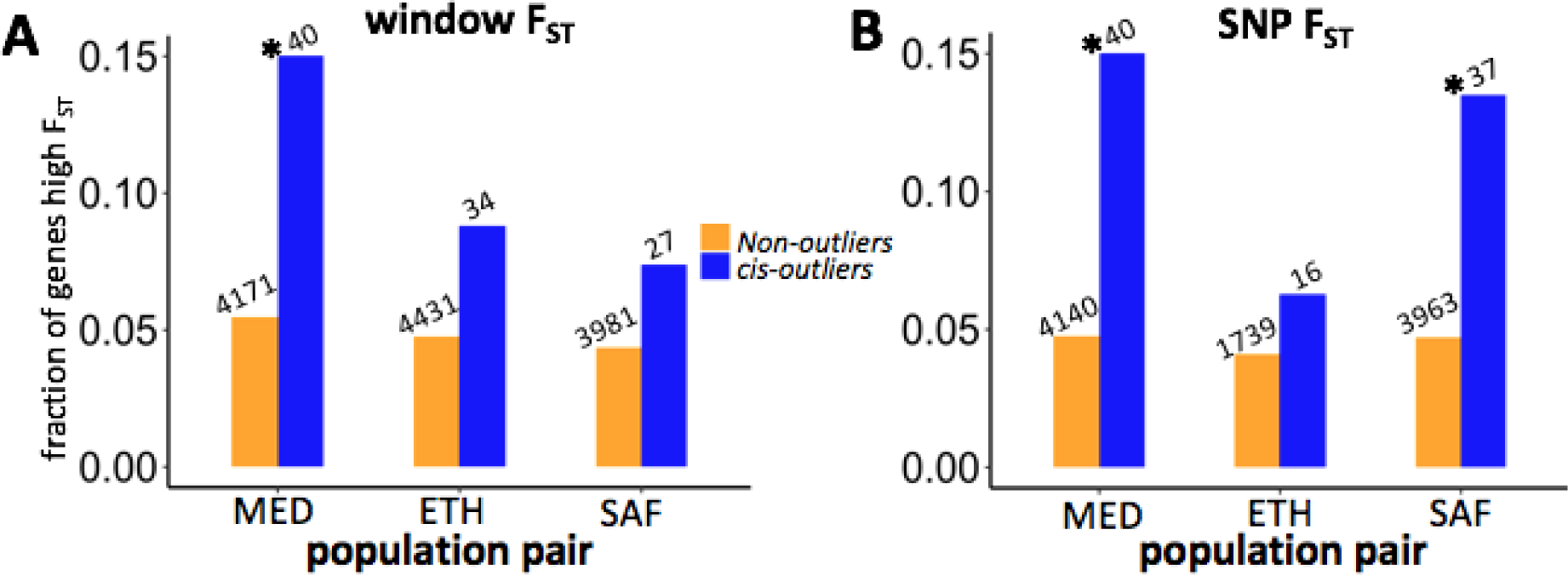
Enrichment of *cis-outliers* for genes with high window *F_ST_* (A) and high SNP *F_ST_* (B) for each population pair. The number above the bar shows the denominator of the fraction. * indicates the fractions are significantly different between *cis*-effect outliers and non-outliers (p < 0.05, Fisher’s Exact Test).

We further examined potential targets of *cis*-regulatory local adaptation, excluding genes with *cis*-effects and trans-effects in the opposite directions. For the *cis*-genes of the MED pair, *Ciao1* and *Cyp6a22* showed high window *F_ST_*, *CG42565* showed high *F_ST_SNPmax_*, and *CG8034* and *Cyp6a17* showed both. *CG8034* (a predicted monocarboxylic acid transporter) was also cited above as having parallel *trans*-regulatory evolution between the MED and ETH pairs, and this gene’s *cis*- and *trans*-effects were both upregulated in FR relative to EG in MED. *CG42565* represents a differentially-spliced product of a transcript that alternatively yields an isoform of *CG13510*, which is upregulated in response to cold (Qin et al. 2005). Interestingly, *Cyp6a22* and *Cyp6a17* belong to the cytochrome P450 protein family (*Cyp6a22* is 248 bp upstream of *Cyp6a17*). This region harbors a polymorphic deletion of *Cyp6a17* which is associated with both colder temperature preference (Kang et al. 2011; Chakraborty et al. 2018) and lower insecticide resistance (Duneau et al. 2018). Based on the diagnostic SNPs for *Cyp6a17* (Good et al. 2014), we found the France-enriched allele for *Cyp6a22* is likely linked to the *Cyp6a17* deletion. Likewise, at the population level, the frequency of intact *Cyp6a17* copy is 0.44 in France and 0.95 in Egypt. Hence, adaptive expression differences may sometimes be driven by gene copy number differentiation between populations. For the *cis*-genes of ETH, *RpL24* showed high window *F_ST_* and *KrT95D* showed high *F_ST_SNPmax_*. For SAF, *GXIVsPLA2* showed high *F_ST_SNPmax_*, while *AGO2* showed both high window *F_ST_* and high *F_ST_SNPmax_*. *AGO2* is involved with antiviral defense and developmental regulation (Deshpande et al. 2005; Nayak et al. 2010) and was previously found to contain fixed differences between European and African populations (Pool 2015). For the genes showing high *F_ST_SNPmax_* in any pair, we plotted the SNP *F_ST_* along the gene region and nearby 20kb to show the sites that may be the most likely targets of selection (Fig. 8). For *CG42565, CG8034* and *KrT95D*, we observed that the highest *F_ST_* sites were located within the gene regions. While for *GXIVsPLA2*, the highest SNP *F_ST_* was 1758 bp downstream of the gene. For *AGO2*, the highest SNP *F_ST_* was 6359 bp downstream but the third highest SNP *F_ST_* was within the gene. Overall, the genetic differentiations between cold- and warm-derived populations around these candidate genes can be quite local, but the linked signal of natural selection can extend further, and there are often multiple SNPs that could represent plausible targets of local adaptation within and outside a given gene region.

**Fig 8.**
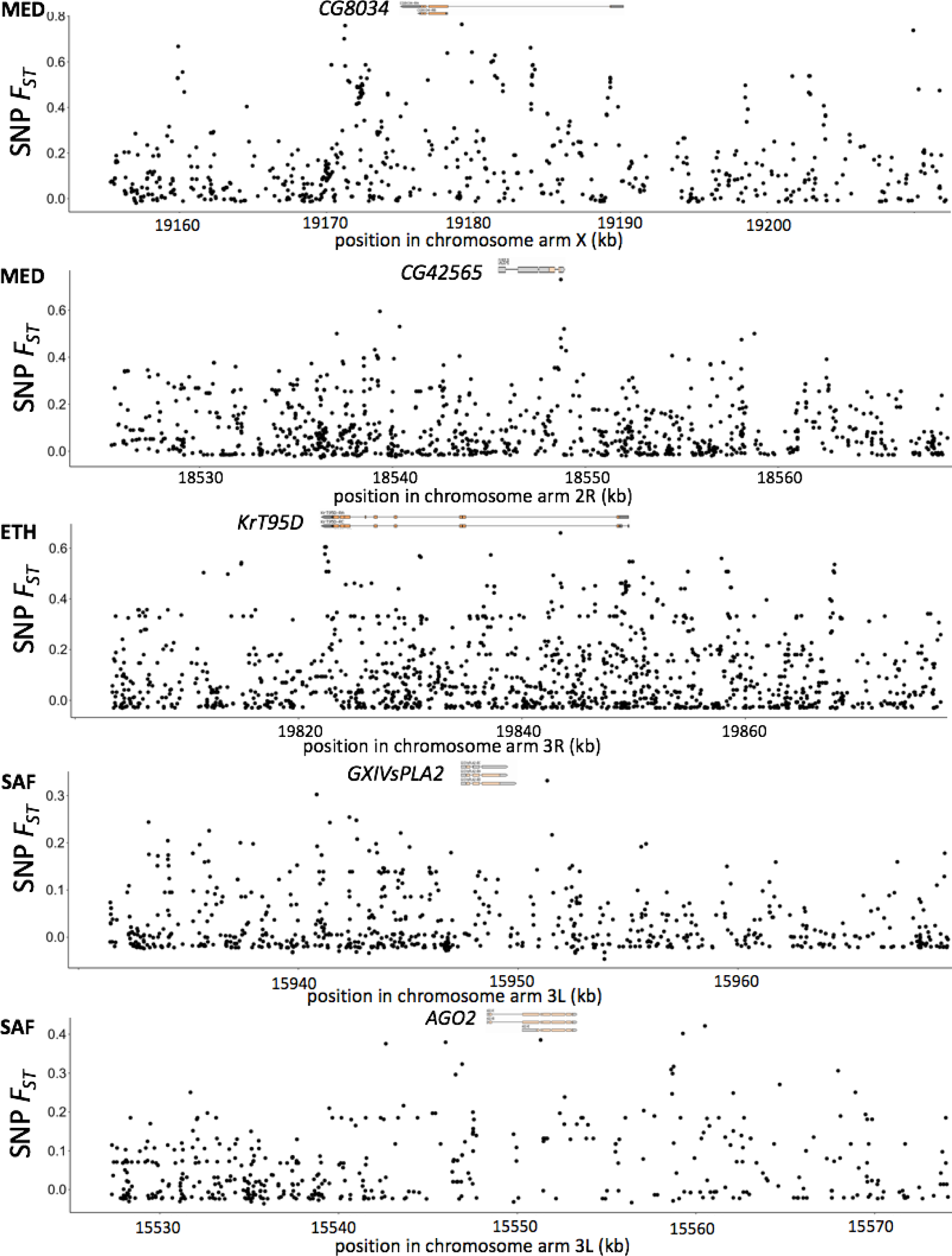
Local peaks of *F_ST_* between warm- and cold-derived populations are observed at some candidate genes for adaptive cis-regulatory evolution. SNP *F_ST_* along the candidate genes with flanking regions of 20kb is shown. The top diagram depicts the coding (orange) and non-coding (gray) exon, captured from GBrowse 2 of *D. melanogaster* (R5.57) from FlyBase (St. Pierre et al. 2014). *Cyp6a17* is not plotted because of the known gene-scale polymorphic deletion.

## Discussion

Parallel evolution has often been studied at the population genetic and trait levels, but it has less frequently been analyzed at the transcriptome level (Stern 2013; Juneja et al. 2016). In this study, we used three recent instances of adaptation to colder climates in *Drosophila melanogaster* to study the evolution of gene expression and alternative splicing. We found a unique pattern of transcriptomic evolution in the high altitude EF sample, involving elevated expression of highly expressed muscle proteins and many other genes. We found the locations of differentially expressed genes on the X chromosome versus the autosomes varies among developmental stages, with the adult female stage having relatively more differentially expressed X-linked genes than the larval stage. We then saw signals of parallel evolution in expression that were higher for larval and adult stages than for pupa. Further, we studied *cis-* and *trans-*regulatory evolution in the context of this ecological adaptation, finding that the relative roles of these regulatory mechanisms differ strongly among population pairs. And we found a signal of *trans-*regulation contributing more predictably to the parallel evolution between population pairs. Finally, those outliers showing *cis*-effects were enriched for high genetic differentiation between populations, suggesting that some of them were the direct targets of selection in cold environments.

Previous comparative transcriptomic studies on multiple pairs of *Drosophila* populations/species have found parallel gene expression differences between high and low latitude populations (Zhao et al. 2015; Juneja et al. 2016). However, because both clines in Australia and North America came from admixtures between European and African ancestry and tropical populations in both clines have a greater proportion of African ancestry (Bergland et al. 2016), it is hard to disentangle adaptive divergence between high and low latitude populations from demographic effects. The demographic influences may be smaller when comparing clines between species pairs (Zhao et al. 2015). Interestingly, the latter study found 10% to 20% differential expressed genes between high and low latitude populations were shared and changed co-directionally between two species *D. melanogaster* and *D. simulans*. The percentages are even higher than what we found between population pairs at adult stages (3% to 10%).

One important reason for the milder patterns of parallelism among our population pairs may be the different selection agents in their unique habitats. In Zhao et al. 2015, samples from the high and low latitude populations from both species were collected in the same areas. While for us, the cold-derived FR population from the MED pair colonized a higher latitude environment than the related warm population, whereas the other two cold-derived populations colonized higher altitude environments where the selection agents may include air pressure, desiccation and ultraviolet radiation (Pool et al. 2017). Although EF and SD have both adapted to higher altitudes (EF at 3,070 meters above sea level, SD at 2,000), SD is seasonally cold (like FR) whereas EF is perpetually cool. We should also note that the standing variation available for adaptation may have differed between our cold-derived populations due to their distinct demographic histories, including the trans-Saharan bottleneck affecting the MED pair and a milder bottleneck in the history of the ETH pair, but no meaningful bottleneck involving the SAF pair (Sprengelmeyer et al. 2020). Hence, although we do significant evidence of parallel gene regulatory adaptation, we suggest that there are both ecological and population genetic reasons to expect substantial non-parallel adaptation as well.

Notably, the EF population exhibits distinct phenotypic evolution such as darker pigmentation (Bastide et al. 2014), larger body size (Pitchers et al. 2013; Lack et al. 2016b), and reduced reproductive rate (Lack et al. 2016b). This distinct evolution in the EF population may explain the strong directionality in gene expression changes between EF and EA for the outliers (Fig. 2), the greater abundance of *trans*-regulatory outliers, and the elevated levels of co-regulation among outliers (Fig. 6). Therefore, the underlying transcriptomic evolution for EF may partly reflect its unique phenotypic evolution, not just adaptation to lower temperature. For example, the upregulation of muscle proteins could reflect the differential abundance of tissues between these Ethiopian populations that differ in size. This size evolution may have also altered the relative proportions of different cell types, which may have driven some of the population differences in gene regulation observed from our whole-organism samples. Future tissue-specific or cell type-specific expression studies involving the EF population can help to examine these possibilities.

We found some evidence of parallel expression evolution between our cold-adapted populations. Developmental stage has a strong effect on the levels of this parallelism, with adult and larva showing significant parallelism while pupa showed a much weaker pattern (Table 1; Figure 4). This is consistent with the observations that larval and adult stages show local adaptation to native temperature but not the pupal stage (Austin and Moehring 2020). It is possible that pupal metamorphosis might reflect a relatively constrained developmental program that limits opportunities for thermal adaptation. The high level of detected parallelism in larvae could also reflect a higher detection power due to less tissue diversity and hence broader spatiotemporal expression of relevant differences. The intriguing pattern of anti-parallelism for some combinations of the pupal stages might suggest that other selective agent is more important than cold (e.g., oxygen level, ultraviolet radiation) for certain population pairs and the direction of selection on gene expression is opposite to the cold. Further, the anti-directional pattern at the pupal stage could be caused by different rates of development for cold-derived populations relative to the warm-derived ones in different pairs. Evidence for such differences is mixed: rates were found to differ between high and low latitude populations in Australia (James & Partridge 1995) but not between our EF and ZI populations (Lack et al. 2016b). Because tissues at different days can generate a wide range of gene expression difference (Hsu et al. 2019), if the cold-derived population develops faster than the warm-derived one in one pair but in another pair the cold-derived population develops slower than the respective warm-derived one, many of the expression differences will be anti-directional between the two pairs. Moreover, because the pupal and larval samples were mixed sex, different rates of development for males and females could led to a biased sex ratio in a sample, especially for pupae (Testa et al. 2013). If the sex ratio bias happened in the cold-derived population in one pair but the warm-derived population in another, it could conceivably result in anti-directional patterns for sex-biased genes.

Compared to the expression abundance, the pattern of parallelism is much weaker for intron usage (Fig S7), which may partly stem from lower power to detect intron usage change (only a small proportion of reads are informative for exon junctions). However, we still found the MED pair and SAF pair show more parallel changes than the combinations with the ETH pair, which is consistent with results for expression abundance. Given the increasing evidence for alternative splicing contributing to environmental response and adaptation (e.g., Singh et al. 2017; Signor and Nuzhdin 2018; Smith et al. 2018), we need to study both expression abundance and splicing to fully understand the evolution at the transcriptome level. The development of sequencing approaches with long reads that cover the entire transcripts (e.g., Iso-Seq) will enable us to quantify isoforms frequency directly and broaden the scope of alternative splicing variation that can readily be quantified. Since splicing changes during development and among tissues (Brown et al. 2014; Gibilisco et al. 2016), a detailed sampling throughout development of different tissues will also be necessary to understand the role of splicing on ecological adaptation.

We found *trans-*effects on expression were more common than the *cis*-effects across the transcriptome (Table 3), which is consistent with some previous studies (e.g., McManus et al. 2010; Coolon et al. 2014; Albert et al. 2018; Glaser-Shmitt et al. 2018) but not with others (e.g., Lemmon et al. 2014; Mack et al. 2016). The transcriptome-wide prevalence of *trans-*effects may be caused by random regulatory changes biased toward *trans*-regulation because of the larger trans-mutational target size (Landry et al. 2007; Metzger et al. 2016). Or, *trans*-regulatory changes may have higher potential for coordinate regulation of multiple genes in networks (Metzger et al. 2016; Liu et al. 2019). To focus on the evolved changes potentially related to adaptation, we compared the proportion of genes with *cis-/trans-*effects for *P_ST_* outliers and to those for non-outliers and saw both effects were enriched in outliers in certain pairs (Fig. 6A & B). These results indicate that the mechanisms of adaptive gene regulatory evolution are highly population-specific, and that either regulatory mechanism has the potential to play a disproportionate role in ecological adaptation.

Moreover, we found a predominance of *trans-*effects associated with parallel outliers than the non-parallel outliers (Fig. 6E & F). In part because of the larger mutational target size of *trans-*regulatory variation for a given target gene, the standing genetic variation for *trans-*regulatory variants may be higher than the *cis-*ones in the ancestral population and therefore the *trans* variants can respond to selection in different population pairs. However, studies in *Arabidopsis thaliana* and *Capsella grandiflora* find that *trans-*eQTLs tend to have lower minor allele frequencies than *cis-*eQTLs (Zhang et al. 2011; Josephs et al. 2020) but it is unclear whether these populations represent the ancestral state before experiencing environmental changes. Also, the potential capacity of *trans-* regulatory factors to co-regulate many genes may amplify the probability of parallel changes between population pairs. Furthermore, since we used whole-body adult samples, it is possible that some *trans-*acting factors regulated genes similarly across tissues while the some *cis-*effects were tissue-specific and were undetected in our mixed-tissue samples. Finally, we emphasize that our study focuses on regulatory changes that may have relatively larger effects (in focusing on *P_ST_* outliers, and in basing *cis*/*trans* analysis on strains showing clearer differences); but small changes may be important for regulatory evolution as well, and may be differentially represented between categories (e.g. *cis* vs. *trans*, parallel vs. non-parallel).

When we considered genes/introns showing both *cis-* and *trans-*effects, we observed that the two types of effects were generally in opposite directions (anti-directional; Table 3). This is consistent with the idea that gene regulation is under stabilizing selection in general and gene regulatory networks evolve negative feedback to buffer effects of regulatory changes (Denby et al. 2012; Coolon et al. 2014; Bader et al. 2015; Fear et al. 2016). With regard to our *P_ST_* outliers, it is possible that *cis-*acting changes might have evolved to compensate for unfavorable pleiotropic impacts of adaptive *trans-*regulatory evolution (or possibly vice-versa). However, negative correlations between *cis-* and *trans-*effects can also be an artifact coming from the measurement error on F1 expression data. Because the F1 data was used to estimate ASE and compared it to 0.5 (*cis-*effect null) and to parental expression proportion (*trans-*effect null), measurement error will introduce artifactual negative correlation between *cis-* and *trans-*acting changes. Therefore, whether the opposing effects between *cis-* and *trans-*acting changes are biologically meaningful will require further study. As Fraser (2019) and Zhang and Emerson (2019) proposed, using independent F1 replicates or other approaches such as eQTL mapping to infer *cis-* and *trans-*effects separately is necessary to affirm evidence of compensatory evolution.

We expect that the adaptive expression divergence caused by *cis-*regulatory changes should leave a signal in the genetic variation of the nearby genomic region. Therefore, we used *F_ST_* statistics to quantify genetic differentiation for the region around the focal genes. Window *F_ST_* is sensitive to classic hard sweeps, and relatively useful for incomplete sweeps and moderately soft sweeps, but it is less useful for soft sweeps with higher initial frequencies of the beneficial allele (Lange and Pool 2016), for which SNP *F_ST_* may be more sensitive. Indeed, a previous genomic study on these same populations found a stronger signal of parallel change for SNP *F_ST_* than for window *F_ST_* genome-wide (Pool et al. 2017). Here, we found genes with outlier *cis-*effects are enriched for those that show high *F_ST_*, especially those with high SNP *F_ST_* (Fig. 7A & B). Hence, standing genetic variants may have contributed importantly to the *cis-*regulatory changes for adaptation in our populations. Genes with both significant *cis-*effects and high *F_ST_* are likely to be the direct targets of the environmental selection and good candidate for future mechanistic studies.

Using three natural fly population pairs with recent adaptive divergence, our study found intriguing patterns of parallel evolution in gene expression and provided new insights on the underlying regulatory effect. In the future, using other approaches to study *cis-* and *trans-*effects in these populations would be necessary, such as eQTL mapping, which can provide more genetic information about the *trans-*regulatory loci. Also, studying the gene expression in different tissues, including different organs from males and females, would provide us a clearer and more comprehensive picture about parallel gene expression evolution. It would also be informative to study other phenotypes besides gene expression that are more related to thermotolerance such as the metabolic pathways identified in this study or nervous system function related to chill coma. Moreover, studying the phenotypic plasticity at different developmental stages could help to explain the different patterns of parallelism in expression evolution across stages and allow us to better understand the importance of local adaptation versus plasticity in thermotolerance.

## Data Availability

The raw RNAseq reads are available from the Sequence Read Archive (SRA) under accessions SRR14179998-SRR14180176 and BioProject PRJNA720479. The population genomics data is from the *Drosophila* Genome Nexus (Lack et al. 2016; http://www.johnpool.net/genomes.html). Custom R and Perl scripts for the *cis*- and trans-effects simulation and analysis can be found at https://github.com/YuhengHuang87/simulation_cis_trans.

## Acknowledgements

We thank Colin Dewey for helpful discussions and the UW-Madison Center for High Throughput Computing (CHTC) for cluster usage. We thank Jeremy Lange for suggestions on *Cyp6a17* gene analysis. This work was funded by NSF DEB grant 1754745 to JEP and by NIH NIGMS grant F32GM106594 to JBL.

## Notes

### Competing Interest Statement

The authors have declared no competing interest.

